# Geranylgeranyl pyrophosphate (GGPP) is associated with hepatic lipid accumulation and insulin resistance in MAO by prenylating Perilipin4

**DOI:** 10.1101/2023.09.19.558458

**Authors:** Yue Zhao, Shan Jiang, Hong-Yu Nie, Meng-Fei Zhao, Peng Sun, Jing-Zi Zhang, Xiao-Chen Wang, Yi-Ping Tang, Xian-Wen Yuan, Xi-Tai Sun, Xiao-Dong Shan, Jian He, Jiang-Huai Liu, Yan Bi, Lei Fang, Xiao Han, Chao-Jun Li

## Abstract

Metabolically Abnormal Obesity (MAO) is characterized by hepatic steatosis and type 2 diabetes (T2D), in contrast to Metabolically Healthy Obesity (MHO). In this study, we investigated the role of hepatic geranylgeranyl pyrophosphate (GGPP), a metabolite of the mevalonate (MVA) pathway, in regulating the differences in lipid metabolism between MAO and MHO. Our findings revealed that GGPP levels were significantly elevated in individuals with MAO, and deficiency of GGPP in the liver ameliorated the defects associated with MAO. Furthermore, we discovered that the prenylation of the lipid droplet-associated protein Perilipin 4 by GGPP enhances the formation of large lipid droplets, thereby exacerbating hepatic lipid accumulation and insulin resistance. Notably, the inhibitor DGBP, targeting the GGPP synthase Ggpps, effectively attenuated the traits of MAO, offering novel insights into the treatment of this condition.

## INTRODUCTION

Obesity has become a global health concern, affecting more than 500 million adults affected worldwide [1,2]. Although obesity is often accompanied by metabolic dysfunctions and inflammation, such as insulin resistance, type 2 diabetes (T2D), hypertension, and cardiovascular disease [3,4], some individuals with obesity remain metabolically healthy, referred to as metabolically healthy obesity (MHO). In contrast, metabolically abnormal obesity (MAO) is characterized by T2D in obese individuals [5,6]. Since T2D is the key unifying factor distinguishing MHO and MAO, identifying the key regulator responsible for MAO-induced T2D may lead to the development of targeted antidiabetic therapeutics to restore adequate insulin sensitivity.

Increased hepatic triglyceride (TG) content (i.e., NAFLD) is a prominent characteristic of MAO individuals [7-10], and the amount of hepatic TG is directly correlated with the degree of insulin resistance in the liver, skeletal muscle, and adipose tissue [11,12]. The progression of NAFLD involves a shift towards large lipid droplet formation within hepatocytes. This transformation is mediated by various lipid droplet-associated proteins, including perilipins and seipins[13-15], which facilitate the maturation of small lipid droplets into larger, more mature structures.

The mechanistic details of perilipin membrane transfer remain an area of active research. Recent studies propose that specific protein-protein interactions and post-translational modifications, such as phosphorylation and acetylation, play a role in regulating perilipin movement between membranes[14,16,17]. Whether there are other post-translation modification methods that affect the membrane translocation of perilipin remains unclear.

Isoprenoid geranylgeranyl diphosphate (GGPP) is a crucial intermediator in the mevalonate pathway (MVA) for cholesterol, terpene, and terpenoid synthesis, which mainly occurred in the liver [18,19]. GGPP is involved in protein prenylation, a post-translational modification essential for the functioning of many peripheral and signal transduction proteins containing cysteine residues in the CaaX motif, such as SREBP1 and CYB5R3 [20,21].

In this study, we employed a metabonomics-based approach to analyze metabolite changes in MAO and MHO individuals and found that GGPP and its synthetase geranylgeranyl diphosphate synthase (Ggpps) protein expression were significantly increased in the livers of MAO patients and mice. With hepatocyte-specific GGPP deficiency mice generated by *Ggpps* knockout (LKO), we revealed that liver GGPP was related to the development of MAO, hepatic lipid droplet formation, and insulin resistance through Perilipin4 prenylation.

## RESULTS

### GGPP and GGPPS are upregulated in the serum and liver of MAO patients

A cohort of 288 patients (119 men, 169 women; 38 ± 22 years old) who underwent laparoscopic Roux-en-Y gastric bypass surgery was enrolled in this study. Characteristics of the cohort categorized separately by T2D are presented in Table 1. Among these subjects, 136 were classified as MAOs with type 2 diabetes (T2D) (52%) and 152 were classified as MHOs without T2D (48%). Notably, MHO individuals had significantly lower fasting blood glucose, 1-hour blood glucose, and 2-hour blood glucose levels compared to T2D MAO individuals after a 75-g glucose load during the oral glucose tolerance test (OGTT). Furthermore, MHO individuals exhibited significantly lower fasting insulin and homeostasis model assessment of insulin resistance (HOMA-IR) values compared to MAO individuals. Despite similar total body fat content, MHO individuals had lower liver fat, visceral fat, and mean adipocyte size than MAO individuals. Conversely, there was no difference in subcutaneous fat between the two groups. In addition, MHO individuals had significantly lower serum alanine aminotransferase (ALT) and aspartate aminotransferase (AST) levels than MAO individuals. After excluding subjects treated with lipid-lowering medications, MHO individuals also exhibited lower fasting triglycerides (TG), total cholesterol, and low-density lipoprotein (LDL) levels compared to MAO individuals. However, there was no significant difference in high-density lipoprotein (HDL) levels between the two groups (Table 1). Overall, these results indicate that the glucose and insulin response, as well as insulin target organs TG deposition, significantly differ between MHO and MAO individuals.

**Table 1.**
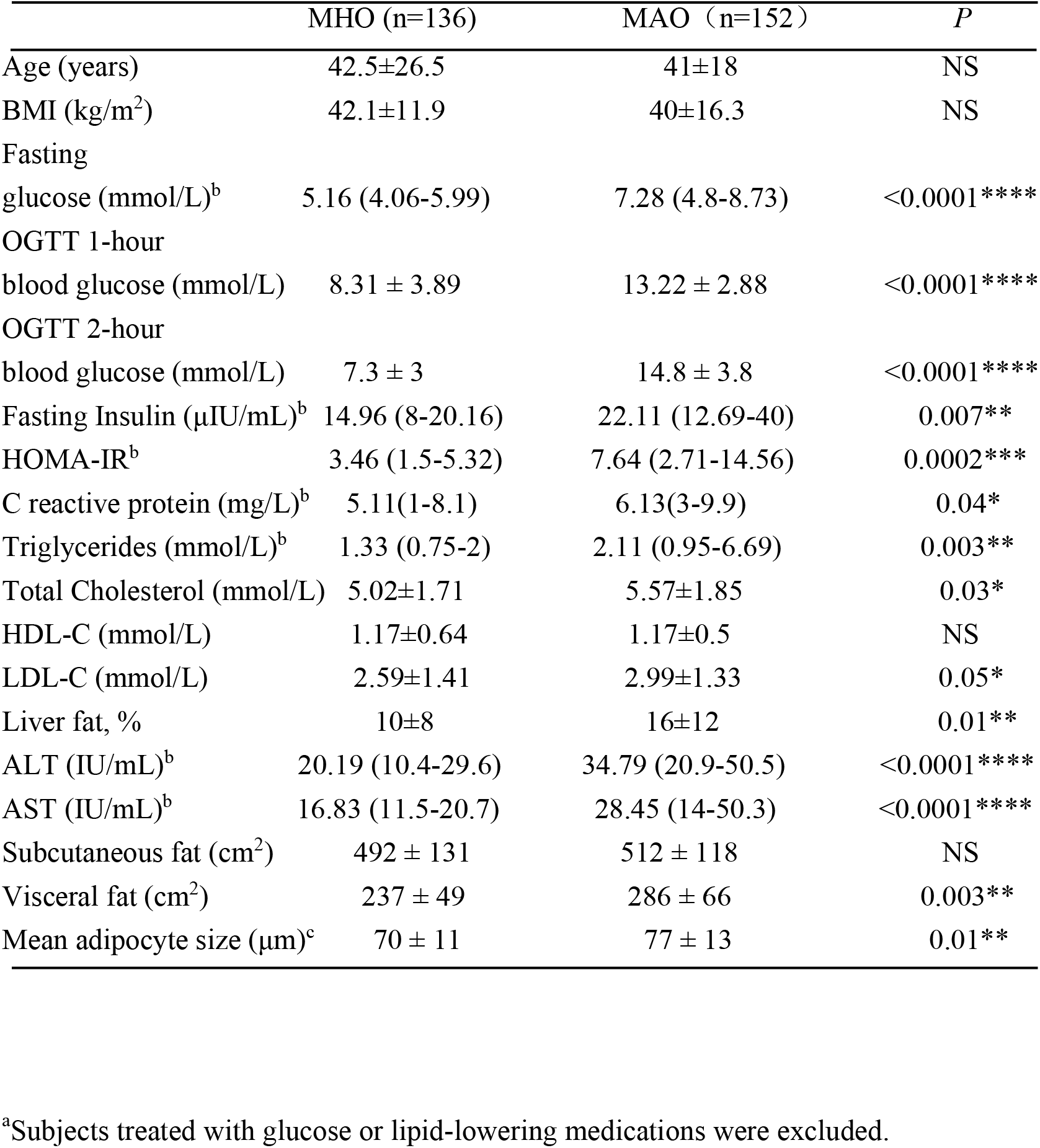
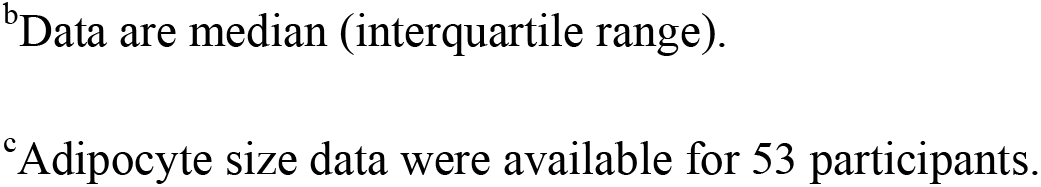
Anthropometric and biochemical parameters of the study subject ^a^.

|  | MHO (n=136) | MAO (n=152) | P |
| --- | --- | --- | --- |
| Age (years) | 42.5±26.5 | 41±18 | NS |
| BMI (kg/m<sup>2</sup>) | 42.1±11.9 | 40±16.3 | NS |
| Fasting |  |  |  |
| glucose (mmol/L)<sup>b</sup> | 5.16 (4.06-5.99) | 7.28 (4.8-8.73) | <0.0001**** |
| OGTT 1-hour |  |  |  |
| blood glucose (mmol/L) | 8.31 ± 3.89 | 13.22 ± 2.88 | <0.0001**** |
| OGTT 2-hour |  |  |  |
| blood glucose (mmol/L) | 7.3 ± 3 | 14.8 ± 3.8 | <0.0001**** |
| Fasting Insulin (μIU/mL)<sup>b</sup> | 14.96 (8-20.16) | 22.11 (12.69-40) | 0.007** |
| HOMA-IR<sup>b</sup> | 3.46 (1.5-5.32) | 7.64 (2.71-14.56) | 0.0002*** |
| C reactive protein (mg/L)<sup>b</sup> | 5.11(1-8.1) | 6.13(3-9.9) | 0.04* |
| Triglycerides (mmol/L)<sup>b</sup> | 1.33 (0.75-2) | 2.11 (0.95-6.69) | 0.003** |
| Total Cholesterol (mmol/L) | 5.02±1.71 | 5.57±1.85 | 0.03* |
| HDL-C (mmol/L) | 1.17±0.64 | 1.17±0.5 | NS |
| LDL-C (mmol/L) | 2.59±1.41 | 2.99±1.33 | 0.05* |
| Liver fat, % | 10±8 | 16±12 | 0.01** |
| ALT (IU/mL)<sup>b</sup> | 20.19 (10.4-29.6) | 34.79 (20.9-50.5) | <0.0001**** |
| AST (IU/mL)<sup>b</sup> | 16.83 (11.5-20.7) | 28.45 (14-50.3) | <0.0001**** |
| Subcutaneous fat (cm<sup>2</sup>) | 492 ± 131 | 512 ± 118 | NS |
| Visceral fat (cm<sup>2</sup>) | 237 ± 49 | 286 ± 66 | 0.003** |
| Mean adipocyte size (μm)<sup>c</sup> | 70 ± 11 | 77 ± 13 | 0.01** |

<sup>a</sup>Subjects treated with glucose or lipid-lowering medications were excluded.
<sup>b</sup>Data are median (interquartile range).
<sup>c</sup>Adipocyte size data were available for 53 participants.

To identify metabolic factors that may contribute to metabolic comorbidities, we collected serum samples from age-matched MHO and MAO subjects (Table 1) and subjected them to ultra-high-performance liquid chromatography and mass spectrometry (UHPLC-MS) for metabonomics analysis. The subsequent Gene Ontology (GO) analysis showed a significant upregulation of steroid metabolism, fatty acid (FA) beta-oxidation, and biosynthesis of unsaturated fatty acids in MAO compared to MHO (Figure 1A). According to GO analysis and KEGG pathway assignment, we found that metabolites involved in cholesterol and lipid metabolisms, such as those related to steroid biosynthesis and fatty acid biosynthesis, were notably upregulated (Figure 1B). Among the 1174 variants analyzed, we identified 55 annotated metabolites with fold change (FC) <-1.5 or >1.5 and P<0.05 (Figure 1C). To narrow down the dataset, we selected several metabolites involved in cholesterol and lipid metabolism that have been reported to be associated with diabetes, indicating reliable metabonomics. Interestingly, GGPP, a metabolite involved in cholesterol and steroid biosynthesis, was found to be elevated in the serum of MAO individuals compared to MHO individuals (Figure 1D). We hypothesized that the high level of serum GGPP was associated with changes in the mevalonate pathway of the MAO liver. Indeed, liver GGPP (Figure 1E) and its synthetase-GGPPS (Figure 1F) were both increased in liver samples from MAO compared to MHO. We confirmed this finding through immunohistochemistry (IHC) and western blotting analyses (Figure 1G) of the clinical liver samples. Larger LD size and TG accumulation were observed in the liver of MAO patients (Figure 1H), and the number of large LDs (>10-μm in diameter) in MAO patients was approximately six-fold higher than that in MHO patients (Figure 3I). Meanwhile, we observed enhanced lipid droplet (LD) size and LD-associated proteins involved in lipid metabolism, such as Perilipin, Cidea, and Rab5, in the liver of MAO individuals compared to MHO individuals (Figure 1J). Furthermore, hepatic *Ggpps* mRNA expression positively correlates with plasma insulin (Figure S1A), glucose (Figure S1B), plasma cholesterol level (Figure S1C), plasma ApoE level (Figure S1D), plasma ApoB level (Figure S1E), and BMI (Figure S1F).

**Figure 1.**
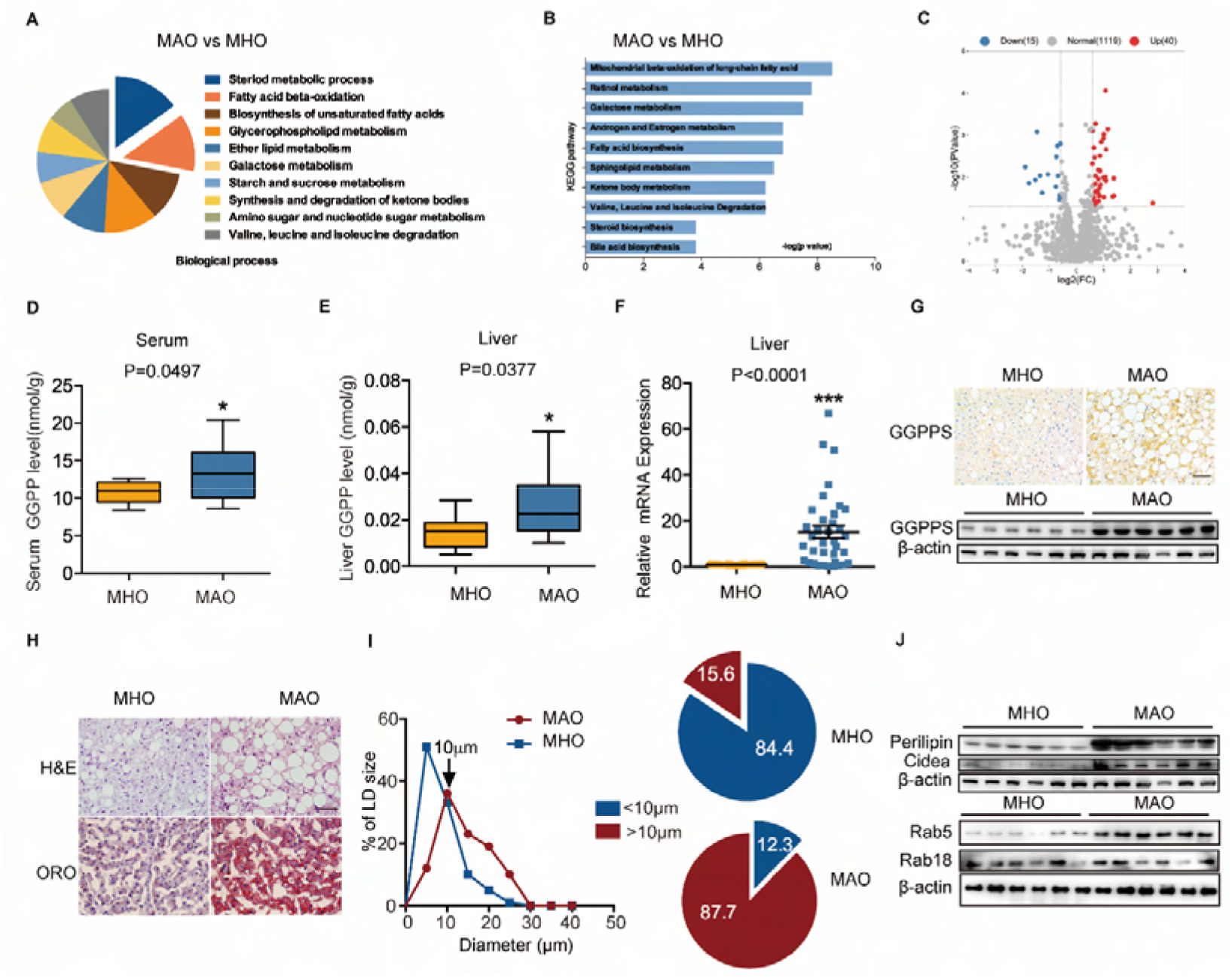
The upregulation of liver GGPP and GGPPS correlates with MAO severity. (A-B) Analysis of biological process (A), KEGG pathway (B) from upregulation of serum metabolites in MAO (n = 10) compared to MHO (n = 10). (C) The volcano plot of differential metabolites in MHO and MAO. The volcano plot was constructed using the fold change values and *P*-adjust. Red dots indicate upregulated genes; blue dots indicate downregulated genes. (D) The Serum GGPP level in MHO (n = 10) and MAO (n = 10). (E-F) The liver GGPP level (n = 10) (E) and relative *Ggpps* mRNA levels (F) of MAO (n = 36) compared with MHO (n = 52). (G) Representative IHC and western blots of GGPPS expression in liver tissues of MHO (n = 6) and MAO (n = 6). (H-I) H&E-staining, Oil Red O staining (H), and quantification of the diameters of lipid droplet (LD) (I) of the liver in MHO and MAO individuals. (J) Western blots of Perilipin4, Cidea, Rab5, Rab18 expression in liver tissues of MHO (n = 6) and MAO (n = 6). All experiments were repeated at least twice with similar results. *p < 0.05, **p < 0.01, ***p < 0.001; *n.s.*, no significant difference. Data are represented as mean ± SEM. Two-sided Student’s t-test. See also Figure S1.

### Liver GGPP accelerates metabolic disorder in MAO animal model

As high levels of GGPP were found in the serum and liver of MAO individuals and MAO mouse models, we generated hepatocyte-specific *Ggpps* LKO mice to explore the effects of hepatic GGPP deficiency on the metabolic change in MAO mice. These LKO mice demonstrated a significant decrease in both Ggpps expression and GGPP level in their livers (Figure S2A). Compared to wild-type mice, hepatocyte-specific GGPP-deficient mice exhibited greater resistance to metabolic disorders. Results from GTT and ITT demonstrated that GGPP deficiency in the liver substantially improved systemic glucose tolerance (Figure 2A) and insulin sensitivity (Figure 2B) in “MAO” mice, resulting in reduced blood glucose and insulin levels (Figure 2C and 2D). To determine the specific tissues involved in insulin sensitivity modulation through hepatic GGPP deficiency, we conducted standard hyperinsulinemia-euglycemic clamp experiments (Figure S2B). GGPP deletion resulted in a slight reduction in glucose levels after constant insulin infusion (Figure 2E). Under clamp conditions, a higher exogenous glucose infusion rate (GIR) was required to maintain the glucose set point in LKO mice (Figure 2F), which was attributed to a lower glucose level because the glucose disposal rate (GDR) was increased (Figure 2G). Although whole-body lipogenesis/glycogen (Figure 2H) was comparable between the two groups, whole-body glycolysis (Figure 2I), liver glucose uptake (Figure 2J), and glycolysis (Figure 2K) were slightly increased. Meanwhile, basal hepatic glucose production (HGP) was significantly suppressed in LKO mice (Figure 2L). Furthermore, glucose uptake by adipose tissue (Figure 2M) and skeletal muscle (Figure 2N) was increased in LKO mice, suggesting that hepatic GGPP may be involved in insulin-mediated metabolic alteration in peripheral tissue.

**Figure 2.**
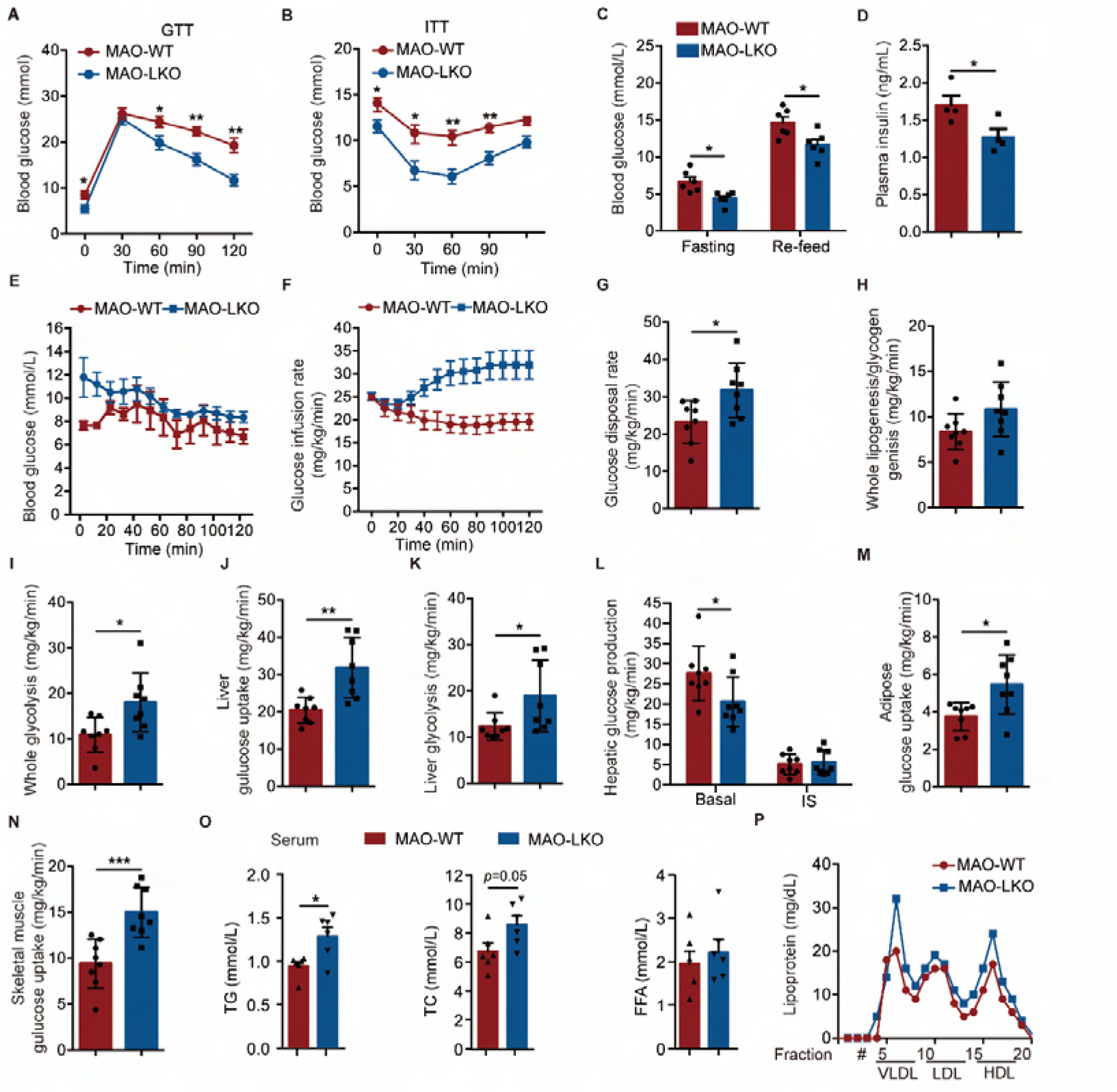
Liver GGPP accelerates MAO defects. (A-B) Glucose (A) and insulin (B) tolerance tests in mice. (n = 8). (C) Blood glucose in overnight-fasted/re-feed MAO-WT and MAO-LKO mice. (n = 6). (D) Plasma insulin in overnight-fasted MAO-WT and MAO-LKO mice. (n = 4). (E-N) Blood glucose (E), Glucose infusion rate (GIR) (F), IS-GDR (G), whole lipogenesis/glycogen genesis (H), whole glycolysis (I), liver glucose uptake (J), liver glycolysis (K) and glucose production (L), adipose glucose uptake (M), and skeletal muscle glucose uptake (N) during hyperinsulinemic-euglycemic clamp studies from MAO-WT (n = 8) and MAO-LKO mice (n = 8). (O) Serum TG, total cholesterol, and free fatty acid (FFA) levels in overnight-fasted WT and LKO mice under MHO or MAO conditions. (n = 6). (P) VLDL, LDL, and HDL transport were detected by FPLC in MAO-WT and MAO-LKO mice. (n = 6). All experiments were repeated at least twice with similar results. *p < 0.05, **p < 0.01, ***p < 0.001. Data are represented as mean ± SEM. Two-sided Student’s t-test. See also Figure S2.

In contrast to the beneficial effect of GGPP deficiency on the liver, plasma TG and total cholesterol levels were increased in *Ggpps*-deficient mice under MAO conditions without changes in free fatty acids (FFA) (Figure 2O). the plasma lipoprotein levels of very-low-density lipoprotein (VLDL), low-density lipoprotein (LDL), and high-density lipoprotein (HDL) were elevated in *Ggpps*-deficient MAO mice (Figure 2P), indicating that liver GGPP may modulate not only the liver but also other endocrine organs to maintain whole metabolic homeostasis by regulating lipid metabolism. Moreover, the large elevation of alanine aminotransferase (ALT) and aspartate aminotransferase (AST) levels in MAO mice was improved after *Ggpps* deletion (Figure S2C). Pyruvate tolerance tests (PTTs) and glucose production assays demonstrated that pyruvate-dependent hepatic glucose output was strongly abrogated in LKO mice (Figure S2D and S2E). Consistently, liver glycogen content and ketone bodies increased (Figure S2F), possibly arising from increased pyruvate production through the glycolytic oxidative pathway, although plasma ketone bodies and lactate levels remained unchanged (Figure S2G). Collectively, our data suggest that GGPP, a metabolite from the MVA pathway in the liver, can regulate glucose and lipid metabolism in insulin-target organs such as the liver and adipose tissue, ultimately accelerating metabolic disorder during MAO progression.

### GGPP elevation is related to LD size and insulin resistance in MAO liver

To validate the clinical and animal findings, primary hepatocytes were treated with oleic acid (OA), which induced a time- and dose-dependent increase in Ggpps expression and GGPP levels (Figure 3A-C). Treatment with GGPP resulted in a significant increase in lipid content in primary hepatocytes (Figure S3A), and knockdown of *Ggpps* led to a significant reduction in TG levels (Figure S3B). Additionally, GGPP treatment led to a significant increase in the average diameter of the LDs in hepatocytes, while GGPP deficiency resulted in the accumulation of smaller LDs (Figure 3D). Furthermore, GGPP treatment impaired the insulin-signaling pathway and insulin-mediated suppression of glucagon-stimulated glucose production in primary hepatocytes (Figure 3E and 3F). GGPP also upregulated lipogenesis gene expression without altering lipolysis genes and inhibited glycolysis gene expression while increasing the expression of gluconeogenesis genes in primary hepatocytes (Figure S3C-F).

**Figure 3.**
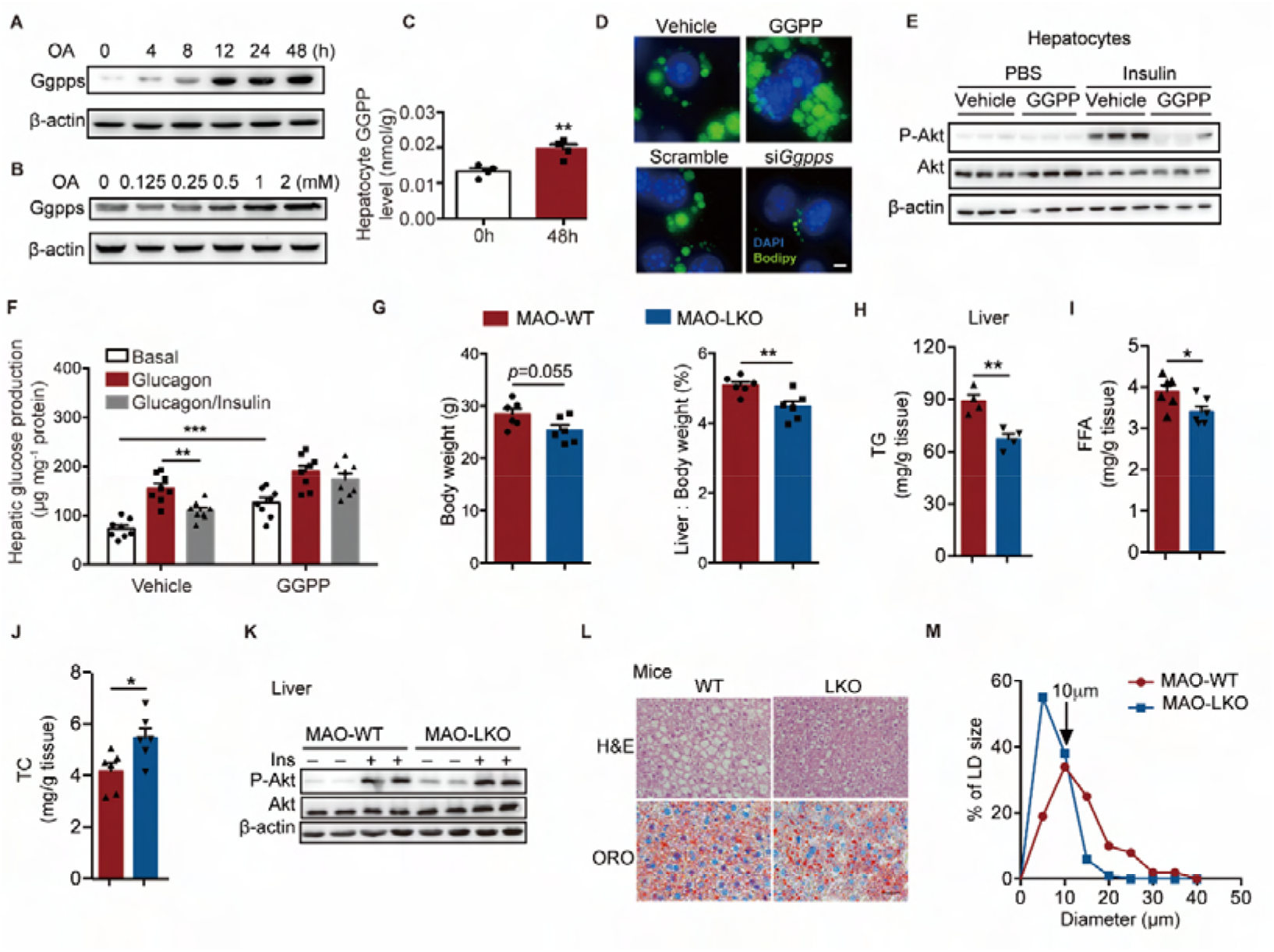
GGPP elevation is related to LD size and insulin resistance in MAO liver. (A) Ggpps protein levels and GGPP levels in primary hepatocytes following the time-dependent OA incubation of primary hepatocytes. (B) Ggpps protein levels in primary hepatocytes following the dose-dependent OA incubation of primary hepatocytes. (C) Primary hepatocyte GGPP level after OA treatment for 48 hours. (D) Fluorescence microscopy of primary hepatocytes treated with vehicle or GGPP and scrambled control (Scramble) or a *Ggpps* siRNA (si*Ggpps*), stained with Bodipy (lipid, green) and DAPI (nucleus, blue). (Scale bar, 5 μm). (E) Insulin-stimulated phosphorylation of AKT in the hepatocytes treated with GGPP. (F) Glucagon-stimulated glucose production in primary hepatocytes within GGPP treatment. (G) Body weight, percentage of the liver weight relative to the whole-body weight of MAO-WT and MAO-LKO mice. (n = 6). (H-J) Liver triglyceride (TG), free fatty acid (FFA), and total cholesterol in overnight-fasted WT and LKO mice under MAO conditions. (n = 6). (K) Insulin-stimulated phosphorylation of AKT in the liver from MAO-WT and MAO-LKO mice. (L-M) H&E-staining, Oil Red O staining (L), and quantification of the diameters of lipid droplet (LD) (M) of the liver in MAO-WT and MAO-LKO mice. Data were collected from H&E-stained sections from three individual mice, five fields per mouse, 10–15 LDs per field in each group, and analyzed using ImageJ software. All experiments were repeated at least twice with similar results. *p < 0.05, **p < 0.01. Data are represented as mean ± SEM. Two-sided Student’s t-test. See also Figure S3.

To further investigate the effect of GGPP on liver function in the context of MAO modeling, we analyzed hepatic lipid metabolism and insulin sensitivity in mice with liver-specific GGPP deficiency. While body weight remained unchanged (Figure 3G), liver weight, TG, and FFA content were reduced in GGPP-deficient mice (Figure 3G-I), whereas total cholesterol in the liver slightly increased (Figure 3J). Hepatic GGPP deficiency consistently enhanced the insulin signaling pathway, as evidenced by increased insulin-stimulated phosphorylation of Akt in the liver (Figure 3K). Notably, liver GGPP deficiency strongly promoted glucose utilization, as indicated by increased mRNA expression of enzymes regulating glycolysis (Figure S3G). In addition, the mRNA expression of key factors promoting gluconeogenesis, i.e., *Pepck* and *Fbp1*, was downregulated in liver GGPP deficiency mice, indicating impaired *de novo* glucose synthesis (Figure S3H).

Moreover, LD diameters measured on ORO-stained sections revealed a significant decrease in the number of LDs larger than 10-μm in the liver of GGPP-deficient mice (Figure 3L and 3M). Taken together, these findings suggest that the elevation of GGPP levels leads to lipid droplet enlargement and insulin resistance in the liver of MAO subjects.

### Liver GGPP regulates lipid droplet formation through prenylation of Perilipin4

Given that GGPP-regulated prenylation is crucial for protein anchored to the membrane surface or embedded in the lipid bilayer [22,23], we hypothesized that GGPP may induce LD formation by prenylating LD-associated proteins to affect LD-mediated lipid homeostasis. To this end, several LD-associated proteins for potential prenylated sites (CXXX motif), including ACSL, ApoA, ApoB, ATGL, CGI58, CIDE A/B/C, COX2, DGAT2, HSL, LPCAT1/2, PLIN 1/2/3/4/5, PNPLA3, Rab5, Rab18, and identified Rab5, Rab18, and Perilipin4 as potential candidates for prenylation (Figure S4A-B). Among these candidates, Perilipin4 was highly expressed in the livers of MAO individuals (Figure 4A) and co-expressed with Ggpps during hepatocellular LD maturation (Figure 4B), positively correlating with Ggpps (Figure S4D). We observed that Perilipin4 prenylation was significantly increased in the livers of hepatic steatosis and decreased in LKO mice (Figure 4C).

**Figure 4.**
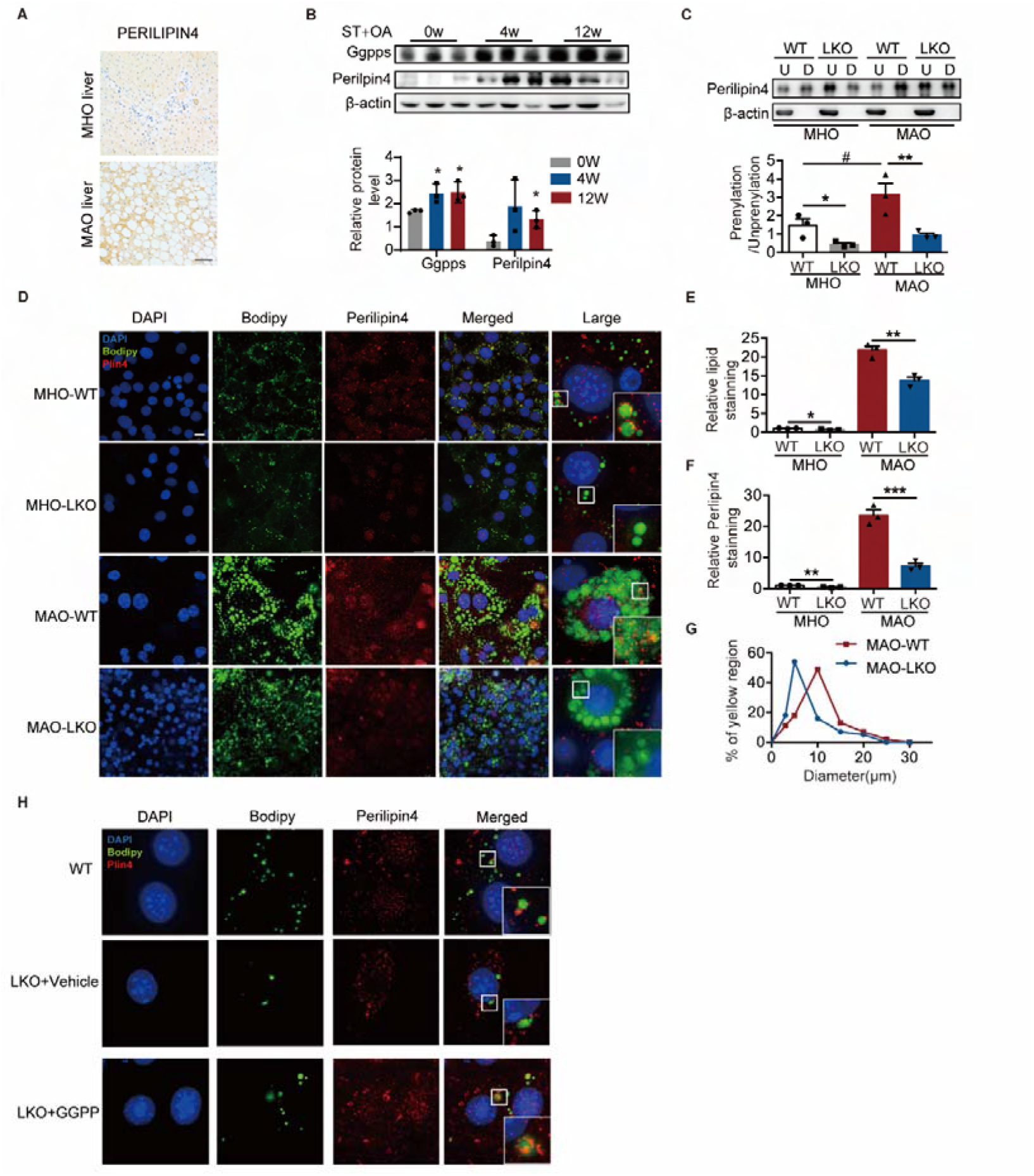
Liver GGPP regulates lipid droplet formation through prenylation of Perilipin4. (A) Representative IHC of PERILIPIN4 expression in liver tissues of MHO and MAO individuals. (B) Ggpps and perilipin4 protein levels in the liver during MAO progression following starch (ST) and oleic (OA) treatment at the indicated times. (C) Membrane-associated Rab5 and Rab18 (lipid-soluble protein with prenylation in the detergent phase, down) and cytoplasm-associated Rab5 and Rab18 (water-soluble protein with no prenylation in the aqueous phase, up) were obtained by Triton X-114 extraction and analyzed by immunoblotting. (n =3). (D-G) Fluorescence microscopy of primary hepatocytes isolated from WT and LKO mice under MHO or MAO conditions, stained with anti-Perilipin4 antibodies (red), Bodipy (lipid, green), and DAPI (nucleus, blue) with quantification of lipid droplet and perilipin4 merged staining. Ggpps and perilipin4 protein levels in the liver during MAO progression following starch (ST) and oleic (OA) treatment at the indicated times. (H) Fluorescence microscopy of primary hepatocytes isolated from LKO mice with or without GGPP, stained with anti-Perilipin4 antibodies (red), Bodipy (lipid, green), and DAPI (nucleus, blue). All experiments were repeated at least twice with similar results. *p < 0.05, **p < 0.01, ***p < 0.001, ^#^p < 0.05. Data are represented as mean ± SEM. Two-sided Student’s t-test. See also Figure S4.

To evaluate the physiological function of GGPP-dependent Perilipin4 prenylation in the liver, we isolated primary hepatocytes in WT and LKO mice to further observe the LD size and lipid accumulation. In WT mice, Perilipin4 normally coated the lipid droplet, whereas Perilipin4 lost the membrane-association function in GGPP-deficient primary hepatocytes as a fluorescence statistic (Figure 4D-G). Furthermore, supplemented with GGPP rescued Perilipin4 membrane-association function in GGPP-deficient hepatocytes (Figure 4H), indicating that Perilipin4 is anchored to the membrane depending on GGPP.

### Plin4C1500S inhibits GGPP-induced hepatic lipid droplet formation and ameliorates GGPP-induced insulin resistance

To demonstrate that GGPP acts by modifying Plin4, we then constructed Plin4C1500S mutant plasmids (geranylgeranylated site mutant) to mimic the un-prenylated state (Figure 5A-B). We observed that cells expressing Perilipin4 mutant showed decreased membrane-association function, and GGPP treatment did not reverse the effects of the Perilipin4 geranylgeranyl site mutation in hepatocytes (Figure 5C lower), suggesting that GGPP regulates hepatic LD formation and lipid accumulation in a Perilipin4 prenylation-dependent manner.

**Figure 5.**
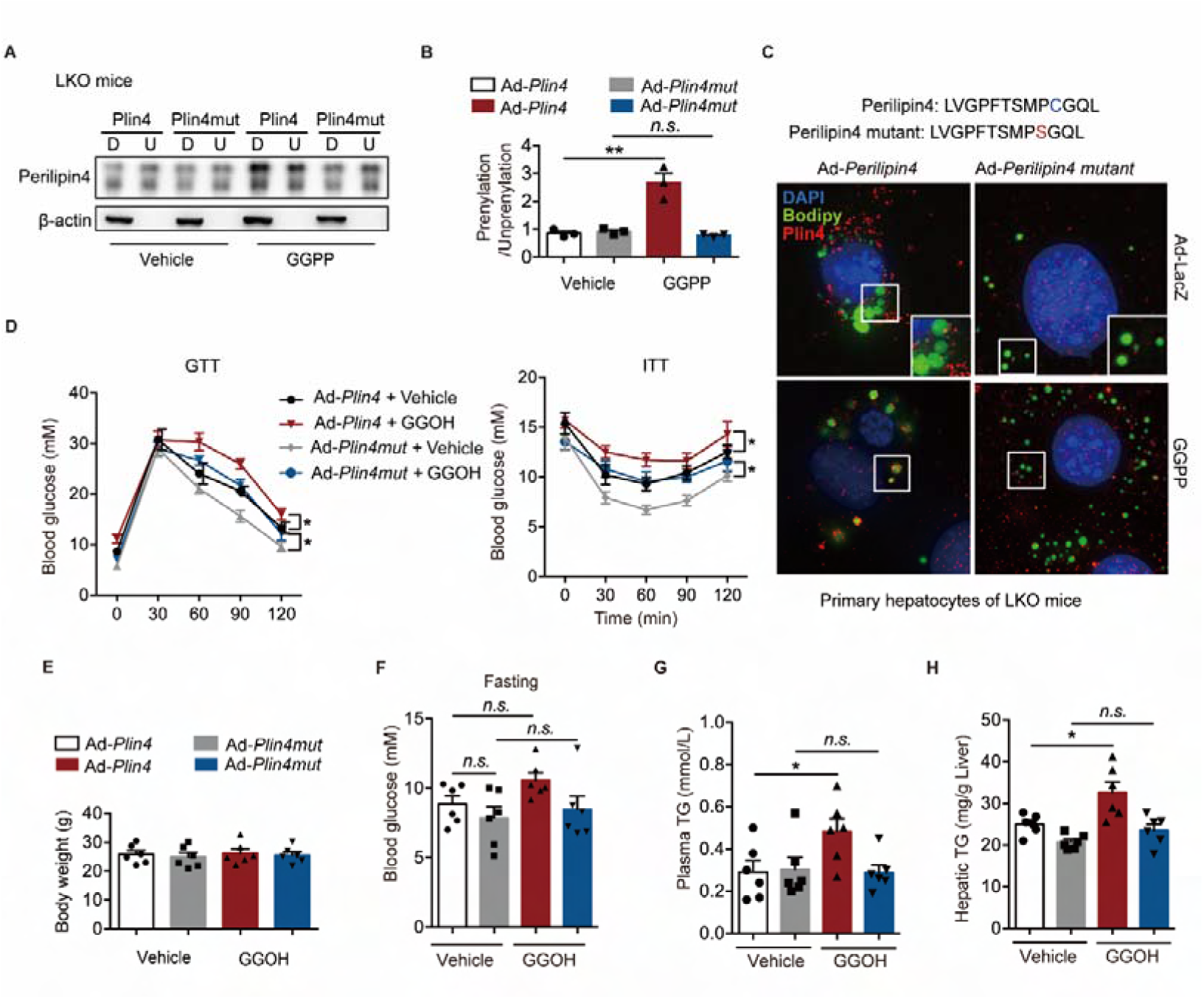
Plin4C1500S inhibits GGPP-induced hepatic lipid droplet formation and ameliorates GGPP-induced insulin resistance. (A-B) Membrane-associated Perilipin4 (lipid-soluble protein with prenylation in the detergent phase, down) and cytoplasm-associated Perilipin4 (water-soluble protein with no prenylation in the aqueous phase, up) were obtained by Triton X-114 extraction and analyzed by immunoblotting in LKO primary hepatocytes treated with vehicle or GGPP and WT or mutant Perilipin4. (C) Fluorescence microscopy of primary hepatocytes treated with vehicle or GGOH and WT or mutant Perilipin4, stained with anti-Perilipin4 antibodies (red), Bodipy (lipid, green), and DAPI (nucleus, blue). (D) Glucose and insulin tolerance tests in mice treated with vehicle or GGPP and WT or mutant Perilipin4 adenovirus. (n = 8). (E) Body weight of LKO mice treated with vehicle or GGOH and WT or mutant Perilipin4. (n = 6). (F-G) Blood glucose and TG in overnight-fasted mice from E. (n = 6). (H) Liver TG in mice from E. (n = 6). All experiments were repeated at least twice with similar results. *p < 0.05, **p < 0.01, ***p < 0.001, *n.s.*, no significant difference. Data are represented as mean ± SEM. Two-sided Student’s t-test. See also Figure S5.

To further investigate the role of GGPP-mediated Perilipin4 prenylation in regulating insulin sensitivity *in vivo*, we compared the systemic glucose tolerance and insulin sensitivity of LKO mice injected with Ad-*Plin4* or Ad-*Plin4* mutant with or without GGOH treatment (GGPP derivative) into the liver.

Systemic glucose tolerance and insulin sensitivity were impaired in the Ad-*Plin4* with GGOH group compared to Ad-*Plin4* with the vehicle group (red v.s. black), the glucose intolerance and insulin resistance still existed between Ad-*Plin4 mutant* with the vehicle and Ad-*Plin4 mutant* with GGOH groups (blue v.s. grey) (Figure 5D). This is most likely due to the presence of endogenous plin4 causing GGPP to still perform some of its functions. The four groups did not differ significantly in body weight (Figure 5E), fasting blood glucose level (Figure 5F), or re-feeding insulin level (Figure S5A). However, blood glucose, TG, and liver TG under the fed condition were higher in the Ad-*Plin4* with GGOH group compared to the Ad-*Plin4* with the vehicle group, but there were few differences between the Ad-*Plin4 mutant* with GGOH and Ad-*Plin4 mutant* with vehicle groups (Figure 5G-H and S5B). These findings indicate that Perilipin4 prenylation affects liver lipid homeostasis. Meanwhile, glucose uptake of iWAT and eWAT unchanged under the Perilipin4 un-prenylation state (Figure S5-D).

### Targeted reduction of hepatic GGPP ameliorates MAO defects

To further define the long-term metabolic effect of hepatic GGPP *in vivo*, we administered GGOH to LKO mice through the tail vein for six weeks. Following treatment, GTTs, and ITTs revealed that GGOH had reversed the improved glucose tolerance and insulin sensitivity observed in LKO mice (Figure 6A). Furthermore, GGOH treatment was able to rescue the abrogation of blood glucose and insulin caused by *Ggpps* deficiency (Figure S6A and S6B), while serum TG levels remained unchanged (Figure S6C). Although body weight and liver weight were comparable to those in the vehicle-treated group (Figure 6B-C), GGOH treatment also reversed the *Ggpps* deletion-induced inhibition of hepatic TG accumulation (Figure 6D and S6D). As expected, GGOH treatment ameliorated the LKO-induced upregulation of hepatic glycolysis (Figure S6E) and downregulation of hepatic gluconeogenesis (Figure S6F). All these data suggest that targeted reduction of hepatic GGPP may ameliorate MAO metabolic defects *in vivo*.

**Figure 6.**
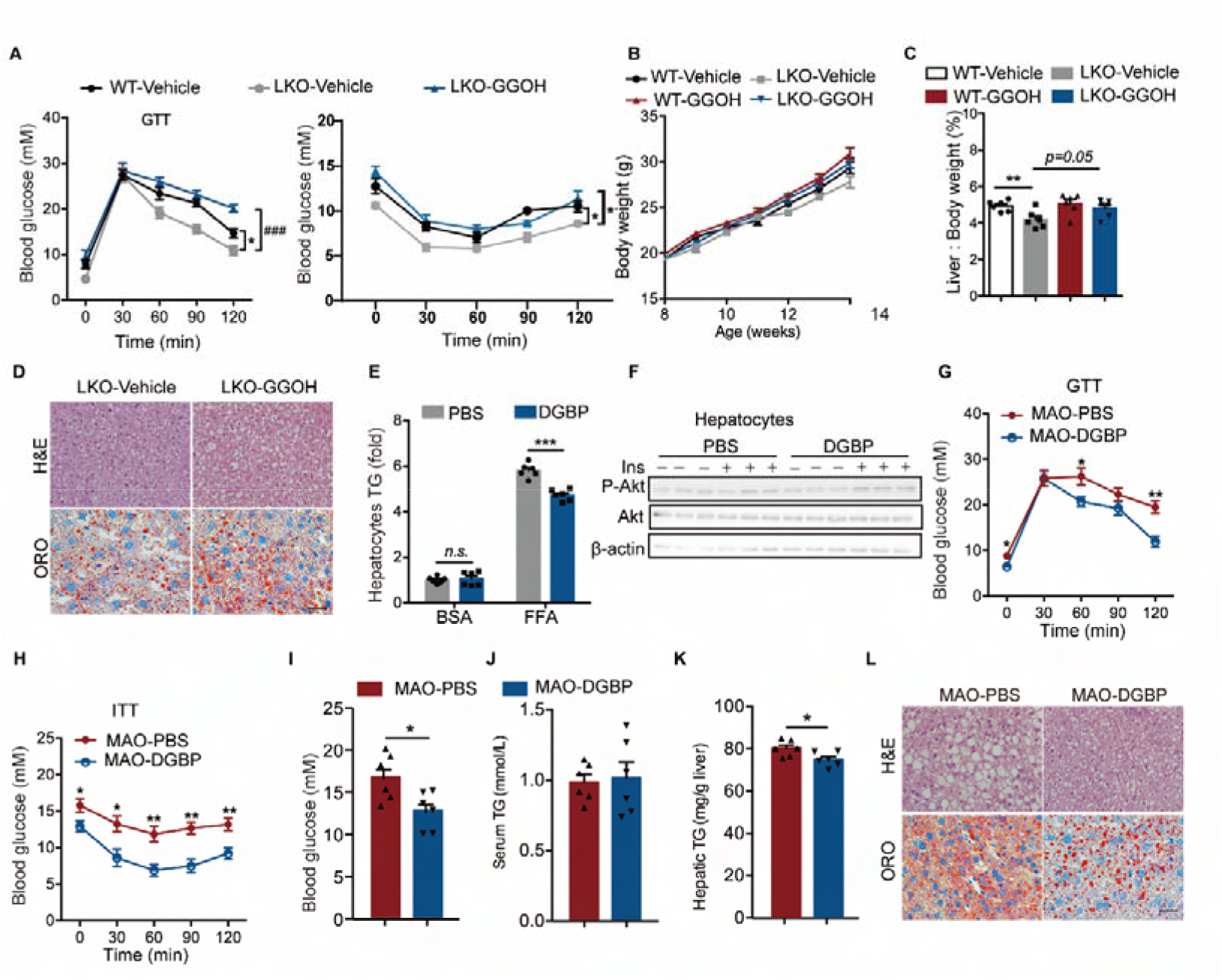
Targeted reduction of hepatic GGPP ameliorates MAO defects. (A) Glucose and insulin tolerance tests of WT or LKO mice with vehicle or GGOH treatment. (n = 8). (B) Body weight of mice from A. (C) Liver weight relative to the whole-body weight of mice from A. (D) H&E-staining and Oil Red O staining of the liver in mice from A. (Scale bar, 50 μm). (E) eWAT weight relative to the whole-body weight of mice from A. (F) Insulin-stimulated phosphorylation of AKT in primary hepatocytes from E. (G-H) Glucose and insulin tolerance tests of MAO mice with PBS or DGBP treatment. (n = 6). (I) Blood glucose in mice from G. (n = 6). (J) Serum TG in mice from G. (n = 6). (N) Hepatic TG in mice from G. (n = 6). (O) H&E-staining and Oil Red O staining of the liver in mice from G. (Scale bar, 50 μm). All experiments were repeated at least twice with similar results. *p < 0.05, **p < 0.01, ***p < 0.001, ^###^p < 0.001. *n.s.*, no significant difference. Data are represented as mean ± SEM. Two-sided Student’s t-test. See also Figure S6.

Therefore, we sought to find a specific inhibitor to interfere with Ggpps for the targeted reduction of hepatic GGPP. DGBP is a potent inhibitor of Ggpps that leads to the depletion of GGPP in the cell [24,25]. We found that incubation of primary hepatocytes with DGBP resulted in enhanced levels of un-prenylated Rap1A (Figure S6H), indicating that GGPP synthesis had been effectively blocked. Furthermore, when hepatocytes were incubated with DGBP and OA, we observed a significant reduction in lipid deposition, as evidenced by the decreased cellular TG levels (Figure 6E). Consistently, insulin signaling was also augmented by DGBP in response to insulin stimulation (Figure 6F). The *in vivo* experiments also show that tail vein injection of DGBP in mice led to improved glucose tolerance and insulin sensitivity (Figure 6G-H), with reduced plasma glucose and insulin concentrations (Figure 6I and S6I), while serum TGs remained unchanged (Figure 6J). Although body weight and liver weight were similar between DGBP-treated and control mice (Figure S6J), hepatic lipid deposition was ameliorated (Figure 6K and 6L). However, we observed impaired overall liver function, as assessed by ALT and AST levels (Figure S6K). Taking these results together, it can be concluded that hepatic GGPP directs obesity classification by modulating hepatic lipid metabolism through covalently prenylating Perilipin4, which underlies the distinction between MAO and MHO. Therefore, targeted reduction of hepatic GGPP can ameliorate MAO defects, inducing the MHO phenotype.

## DISCUSSION

The prevalence of obesity has led to the identification of two distinct phenotypes: metabolically healthy obese (MHO) and metabolically abnormal obese (MAO), based on the presence or absence of metabolic abnormalities such as T2D, hepatic steatosis, and visceral fat accumulation. However, little is known about the mechanisms underlying these differences in metabolic profiles and hepatic lipid accumulation between the two groups. To address this knowledge gap, our study aimed to identify the organs and metabolic pathways that determine the metabolic differences between MAO and MHO. We found that the liver and its mevalonate (MVA) metabolites, specifically geranylgeranyl pyrophosphate (GGPP), play a crucial role in the distinction between MAO and MHO. The metabolites involved in cholesterol and lipid metabolism were notably upregulated in the liver of MAO, and GGPP, an intermediary in the mevalonate pathway for cholesterol synthesis, was found to be responsible for the insulin resistance and hepatic lipid accumulation that distinguishes MHO and MAO. Our results indicate that GGPP regulates hepatic triglyceride accumulation through covalently perilipin4 prenylation-dependent lipid droplet formation.

GGPP, an isoprenoid molecule synthesized by Ggpps in the mevalonate pathway involved in cholesterol synthesis, could play a role in a prenylation-dependent manner or a prenylation-independent manner, respectively. Our previous studies have shown that Ggpps-dependent prenylation is a crucial mediator linking protein prenylation and metabolic reprogramming, causing NAFLD and subsequent fibrosis development [26], lipid-induced muscle insulin resistance [27]. In this study, we highlight the effects of hepatic GGPP on glucose and lipid metabolism in the liver. GGPP can switch a ‘fat but healthy’ insulin-sensitive state to a relatively ‘lipodystrophic’ insulin-resistant animal by prenylating LD-associated proteins Perilipin4.

In current studies, we also found that the chemical DGBP, a potent inhibitor of Ggpps, can suppress MAO defects (Figure 7G-O). Interestingly, FPP, an intermediate metabolite in the mevalonate pathway, has been shown to activate the farnesoid X receptor (FXR), whose agonist-obeticholic acid (OCA) was associated with improved insulin sensitivity compared with placebo (+24.5% vs -5.5%, p=0.011) in a six-week hyperinsulinaemic, euglycaemic clamp study of 64 patients with NAFLD and diabetes [28]. Therefore, we propose that targeting metabolites in the mevalonate pathway may be a viable strategy for treating insulin resistance and hepatic lipid accumulation.

Previous studies identify a large number of biomarkers in the diagnosis of obesity or diabetes; however, none of them have been effectively utilized in the distinction of MAO and MHO [29,30]. Recent studies have revealed that almost 70% of obese individuals exhibit hepatic lipid accumulation, glucose intolerance, and insulin resistance [31], yet the underlying factors driving the severity of MAO are not well understood. In this study, we used a metabonomics analysis to construct a comprehensive database of MHO and MAO. Our findings shed light on the diverse effects of hepatic intermediate metabolite-GGPP in the mevalonate pathway as a key factor in lipid metabolism and hepatic lipid accumulation in liver. Additionally, we have identified GGPP modulates hepatic lipid accumulation via protein covalent modification mechanisms. Given these results, it is plausible that measuring GGPP levels in the liver or serum could serve as a useful biomarker for determining obesity classification, severity, and progression in clinical settings. Further elucidation of these mechanisms could lead to the identification of novel therapeutic targets for the treatment of obesity and related metabolic disorders.

### Limitations of study

Despite our findings, our study has certain limitations that should be considered. Firstly, although we used OGTT to distinguish MAO from MHO, the clamp test could have served as a more precise gold standard for this purpose. Additionally, while we used starch with palmitate or oleate treatment to induce MHO and MAO phenotypes in mice, this model may not fully mimic the clinical conditions observed in humans [32]. Long-term tendencies of mice to gain body weight, liver weight, and adipose weight may differ from those observed in humans. Future studies could address this limitation by using other mouse models of MHO and MAO (such as sucrose with palmitate or oleate) as well as clinical samples with well-defined etiology of MAO. Moreover, while our study focused on the regulation of lipid and glucose metabolism by GGPP, other energy metabolisms in MAO individuals should also be evaluated in future research. Lastly, further studies are needed to investigate the upstream mechanism responsible for the elevated expression of GGPP in MAO. On the other hand, whether the rise of GGPP in the blood could have an effect on other tissues or organs besides the liver needs to be further investigated

## STAR METHODS

### KEY RESOURCES TABLE

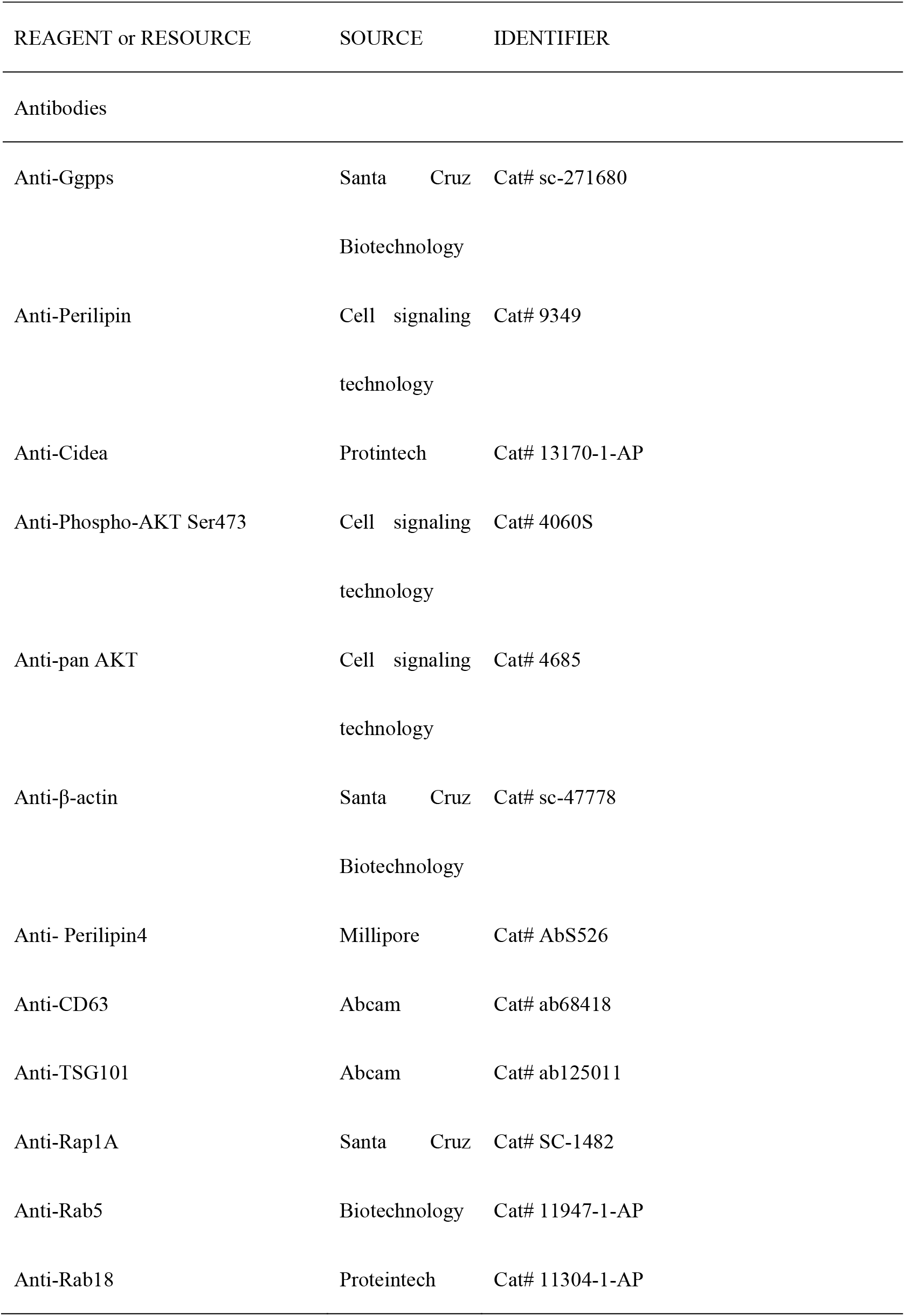
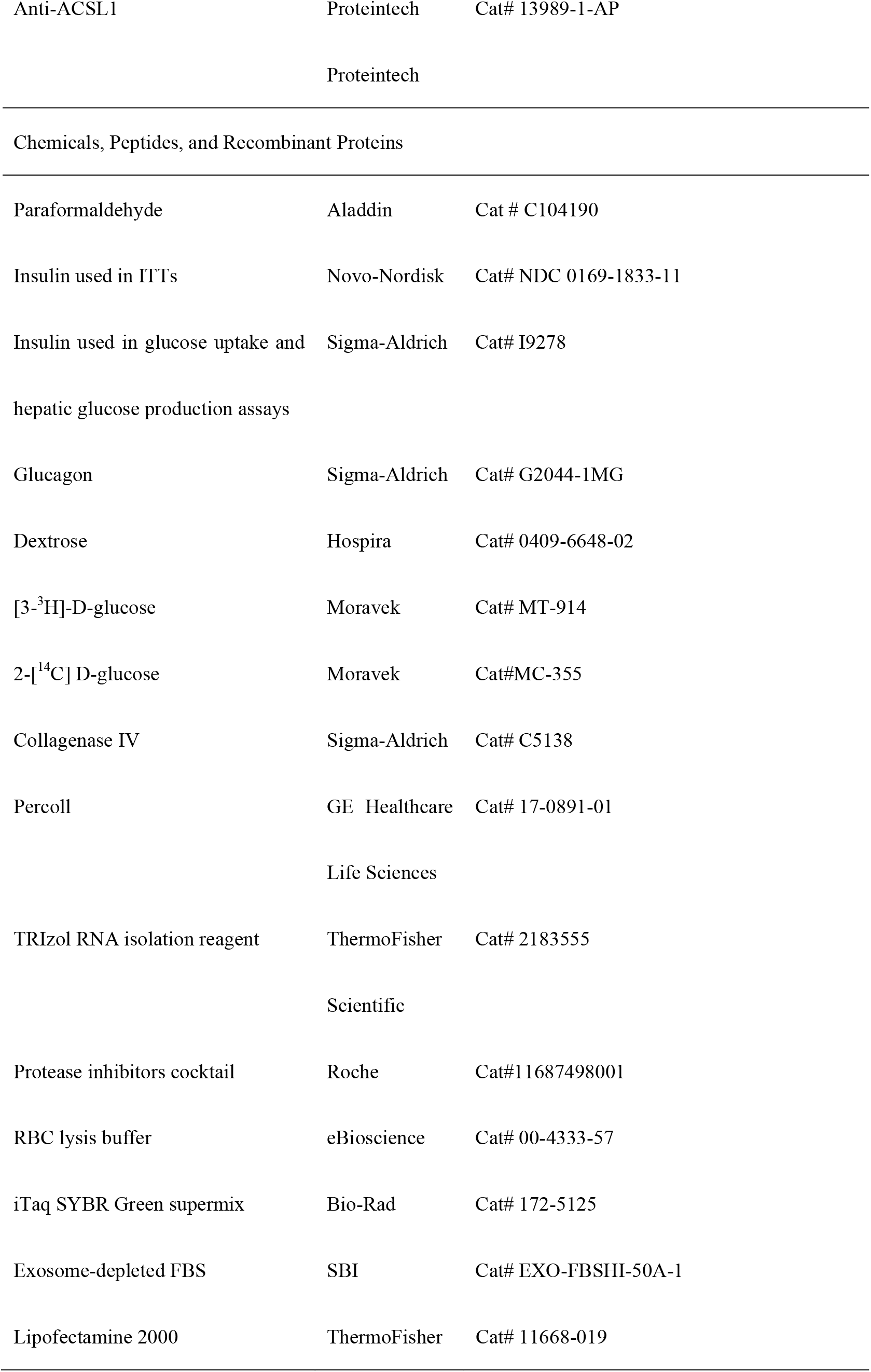
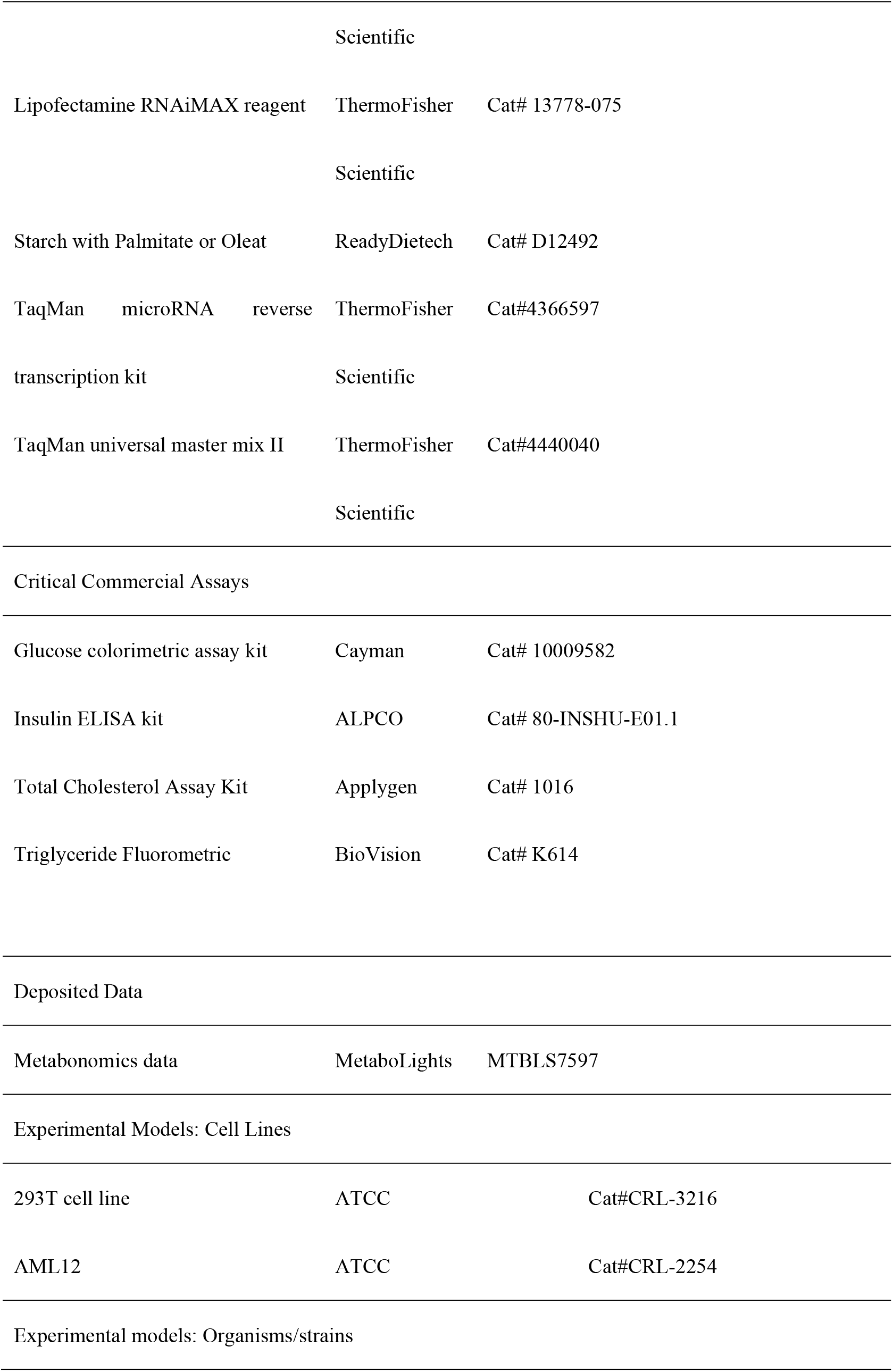
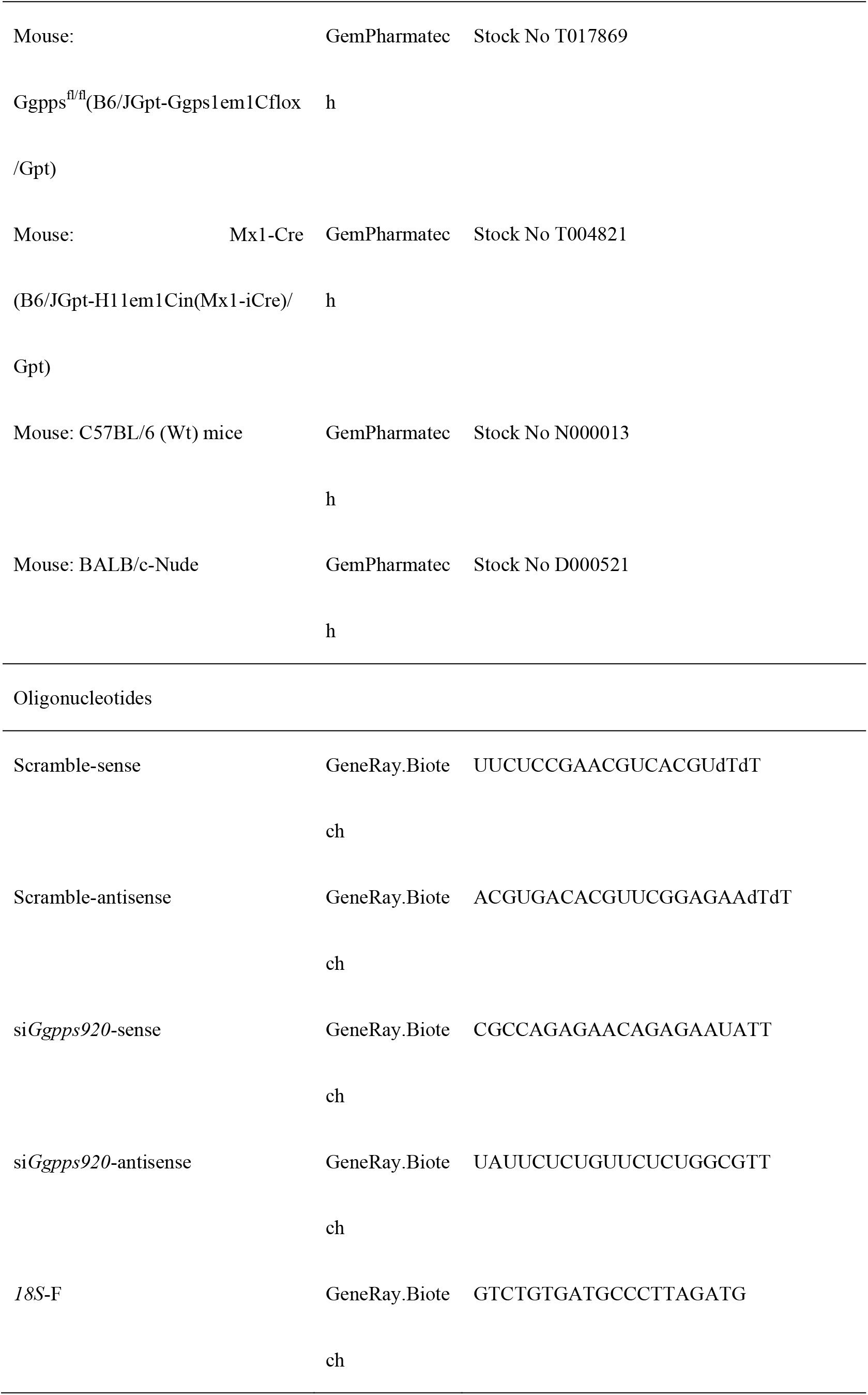
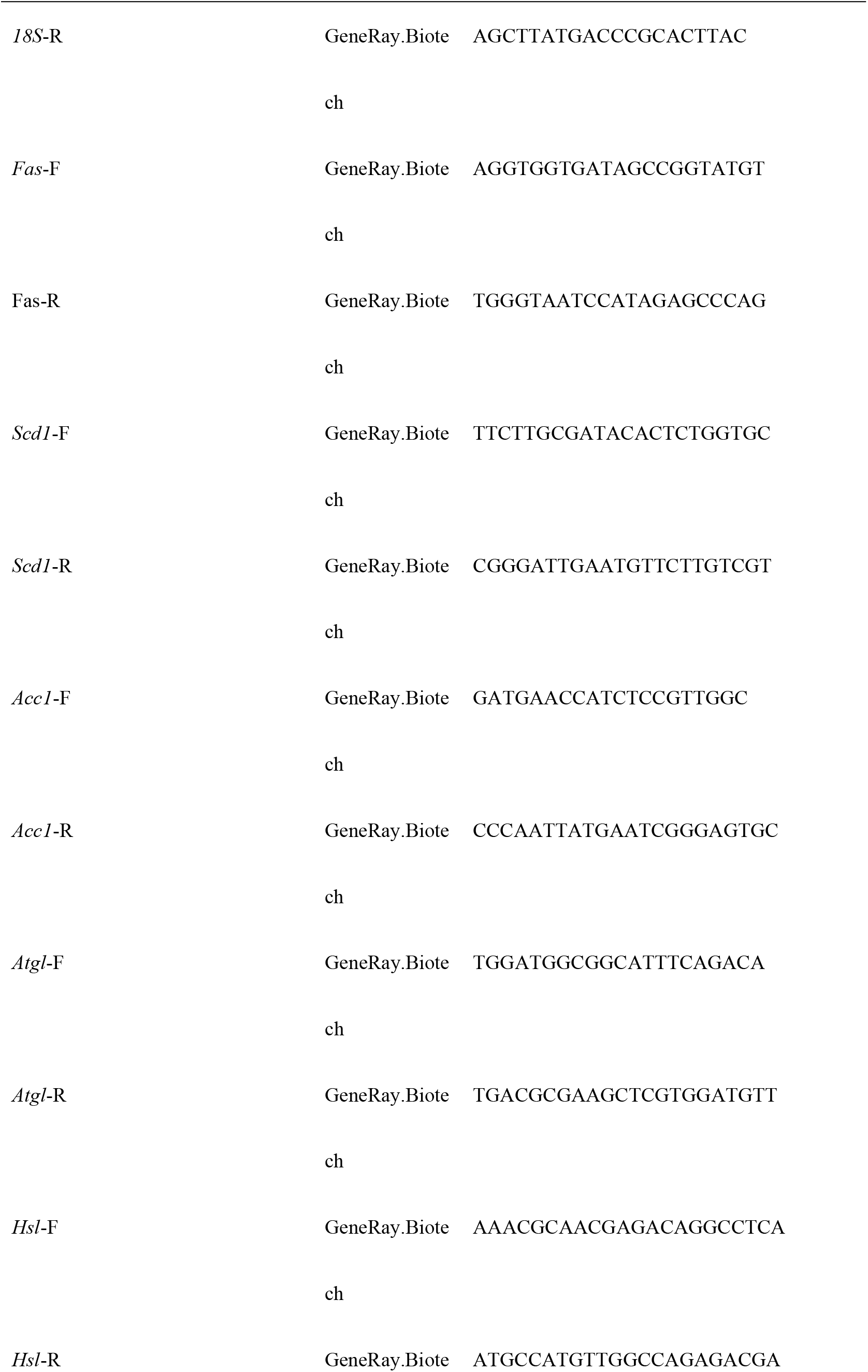
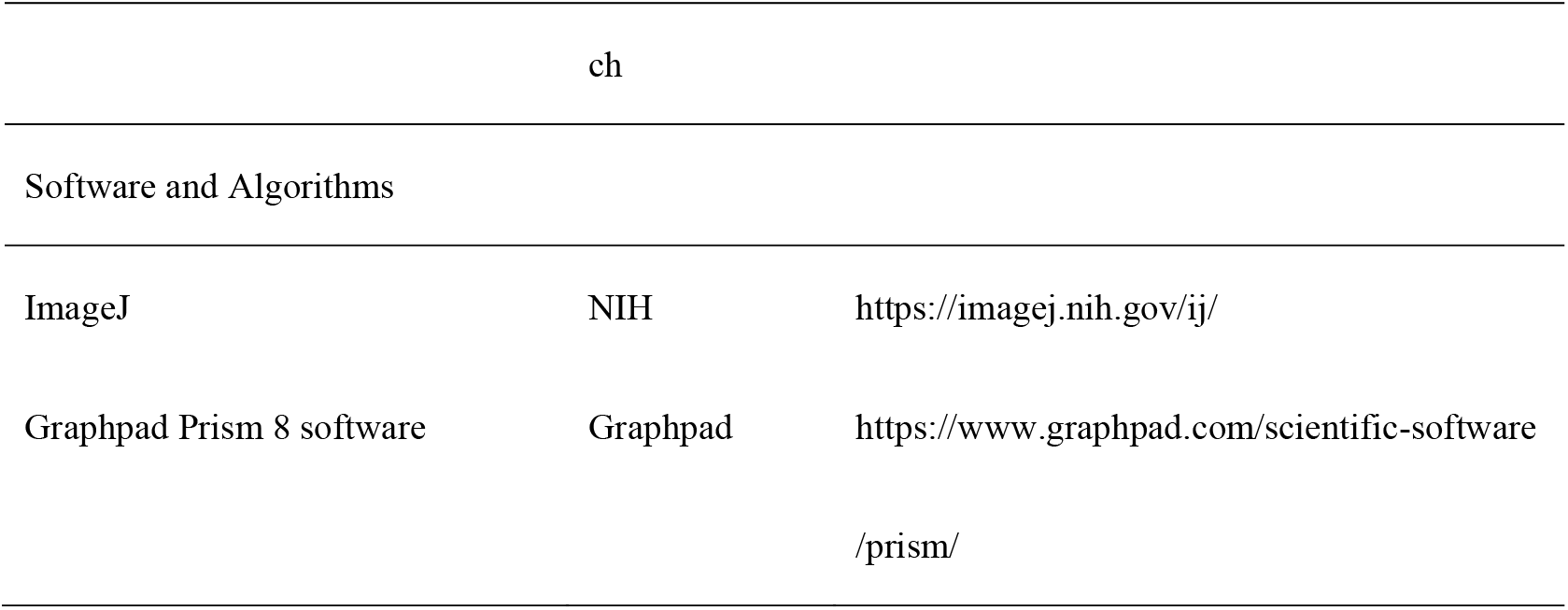

## RESOURCE AVAILABILITY

### Lead contact

Further information and requests for resources and reagents should be directed to and will be fulfilled by the Lead Contact, Chaojun Li.

### Materials availability

All materials used in this study are either commercially available or obtained through collaboration, and are available from the corresponding authors upon request.

### Data and code availability

Metabonomics data are deposited in the MetaboLights: https://www.ebi.ac.uk/metabolights/MTBLS7597.

## EXPERIMENTAL MODEL AND SUBJECT DETAILS

### Mice

C57BL/6J mice were purchased from the Model Animal Research Center of Nanjing University. Liver-specific *Ggpps* knockout mice were generated by crossing *Ggpps*-floxed mice with *Mx1*-Cre transgenic mice. All mice were kept under standard conditions with a 12 hr/12 hr light-dark cycle in temperature 20℃- 22℃ and humidity (50-60 %) controlled cabinets under Specific Pathogen Free (SPF) conditions. The mice were given access to a regular laboratory chow diet (Xietong Bio., Jiangsu, China) and sterilized water ad libitum. For the MHO and MAO model, WT and *Ggpps* LKO mice were fed a starch with palmitate or oleate (ReadyDietech) for 6 weeks starting at 8 weeks of age. *Mx1*-Cre expression can be driven by injection of double-stranded RNA polyinosinic–polycytidylic acid (PIPC), resulting in Cre-mediated recombination confined to the liver. *Ggpps* knockout was achieved by PIPC injection at 7 weeks of age, and the knockout efficiency was detected 1 week later. The animals were given access to water ad libitum. All mice used in the experiments were male. We have complied with all relevant ethical regulations for animal testing and research. All animal procedures were carried out with the approval of the Animal Care and Use Committee at the Model Animal Research Center of Nanjing University in Nanjing, China, and used protocols approved by the institutional animal care committee (AP number: #CS20).

### Human samples

The obese patients in the Endocrinology Department of Drum Tower Hospital Affiliated with the Medical School of Nanjing University were classified into MHO and MAO according to the previous study [6,33]. The patients did not have the following exclusion criteria: (1) use of corticosteroids and glucose or lipid-lowering medications within the last 3 months; (2) acute infection;(3) autoimmune disease; (4) severe cardiovascular and cerebrovascular diseases. Before the surgery, the individuals underwent OGTT and their blood samples were collected. The human samples and data were collected from patients who underwent Roux-en-Y. NAFLD diagnosis was confirmed by liver biopsies in all patients. Those with insulin resistance and/or NAFLD were identified as MAO. The study protocol was approved by the Ethical Committee of the Drum Tower Hospital Affiliated with the Medical School of Nanjing University, and written informed consent was obtained from each participant by the ethical guideline of the 1975 Declaration of Helsinki of the World Medical Association. This study was registered with the International Clinical Trial Registry Platform (ICTRP), with the clinical trial number NCT03296605 as an observational design. The clinical data of all subjects are summarized in Table 1.

### Isolation of primary hepatocytes

Primary hepatocytes were prepared by collagenase digestion via catheterization of the inferior vena cava (IVC) [34]. Before collagenase infusion, the liver was perfused (3–4 min) via IVC after severing the portal vein. When the color of the liver changed to light brown, perfusion was continued with collagenase for 2 min. Within 2 min of digestion, the liver was monitored for the appearance of cracking on the surface. Perfusion was stopped immediately and the liver was excised into an ice-chilled Buffer. Cells were filtered through a 100 μm nylon filter and centrifuged at 60 × g for 6 min at 4℃. The cell pellet was washed once in Buffer (without collagenase) by resuspending and centrifuging at 60 × g for 6 min at 4℃. The cell pellet was then mixed with Percoll and centrifuged at 100 × g for 10 min. Hepatocytes were collected as a pellet and washed once with Buffer (no collagenase) and then cultured in a medium containing penicillin, streptomycin, and 10% FBS. After overnight incubation, the culture medium was switched to EV-free Dulbecco’s Modified Eagle Medium (DMEM) supplemented with antibiotics.

## METHOD DETAILS

### Serum metabolites extraction and UHPLC-MS/MS analysis

The samples (100 μL) were placed in the EP tubes and resuspended with prechilled 80% methanol by well vortex. Then the samples were incubated on ice for 5 min and centrifuged at 15,000 g, 4°C for 20 min. Some of supernatant was diluted to final concentration containing 53% methanol by LC-MS grade water. The samples were subsequently transferred to a fresh Eppendorf tube and then were centrifuged at 15000 g, 4°C for 20 min. Finally, the supernatant was injected into the LC-MS/MS system analysis.

UHPLC-MS/MS analyses were performed using a Vanquish UHPLC system (ThermoFisher, Germany) coupled with an Orbitrap Q ExactiveTM HF mass spectrometer (Thermo Fisher, Germany) in Novogene Co., Ltd. (Beijing, China). Samples were injected onto a Hypersil Goldcolumn (100 × 2.1 mm, 1.9μm) using a 17-min linear gradient at a flow rate of 0.2 mL/min. The eluents for the positive polarity mode were eluent A (0.1% FA in Water) and eluent B (Methanol). The eluents for the negative polarity mode were eluent A (5 mM ammonium acetate, pH 9.0) and eluent B (Methanol). The solvent gradient was set as follows: 2% B, 1.5 min; 2-100% B, 3 min; 100% B, 10 min; 100-2% B, 10.1 min, 2% B, 12 min. QExactive^TM^ HF mass spectrometer was operated in positive/negative polarity mode with a spray voltage of 3.5 kV, a capillary temperature of 320°C, a sheath gas flow rate of 35 psi, and an aux gas flow rate of 10 L/min, an S-lens RF level of 60, Aux gas heater temperature of 350°C.

### GGPP level determination

For sample pretreatment, each 20 mg liver tissue sample was homogenized with 200 μL mixed solvent (isopropanol: 100mM ammonium bicarbonate = 1:1), and then added 800 μL methanol. The mixtures were centrifuged at 18,000 g for 20 min at 4 °C. The supernatant of each sample was transferred to another tube and then dried with nitrogen. The samples were redissolved with 50 μL 50 % methanol and centrifuged for UPLC-MS analysis.

Ultra-performance liquid chromatography coupled to a tandem mass spectrometry (UPLC-MS/MS) system (ACQUITY UPLC H-Class/Xevo G2 TQ-XS MS/MS, Waters Corp., Milford, MA) was used to quantitate GGPP and FPP. All chromatographic separations were performed with an ACQUITY BEH C18 column (1.7 μm, 50 mm × 2.1 mm internal dimensions) (Waters Corp., Milford, MA). The mobile phase consisted of 0.3% ammonia water containing 10 mM ammonium acetate (mobile phase A) and 80 % acetonitrile containing 50 mM ammonium formate and 0.3 % ammonia water (mobile phase B). The column was maintained at 40 °C and the sample manager was at 10 °C. The injection volume for all samples was 5 mL. The flow rate was 0.5 mL/min with the following mobile phase gradient: 0-0.5min (5% B), 0.5–1.5 min (5-30% B), 1.5-4min (30-65% B), 4-4.5 min (65-95% B), 4.5-5.5 min (95%, 5.5-5.7 min (95-5% B), and 6.5 min (5% B). The mass spectrometer was operated in negative ion mode with a 2.0 kV capillary voltage. The source and desolvation gas temperatures were 150 °C and 550 °C, respectively. The desolvation gas flow rate was 1000 L/Hr.

The Raw data generated by UPLC-MS/MS were processed using Masslynx software (Waters Corp., Milford, MA) and QuanMET software (v2.0, Metabo-Profile, Co., Ltd, Shanghai, China) to perform peak integration, calibration, and quantification for each metabolite. Standard curves were generated with 0.5, 1, 5, 10, 20, 50, 80, and 100 nM of GGPP and FPP for absolute quantitation. The linear relationship between the instrument response value and the concentration was calculated to obtain y = ax + b (y represents the instrument response value, such as peak height and peak area).

The mass spectrometry metabolomics data have been deposited to MetaboLights with the dataset identifier MTBLS2909 (www.ebi.ac.uk/metabolights/MTBLS2909) [35].

### Glucose tolerance and insulin tolerance tests

For glucose tolerance tests, mice received one dose of dextrose (1 g/kg body weight) via i.p. injection after 12 h of fasting. Blood glucose concentrations were measured before and 30, 60, 90, and 120 min after the glucose injection. For insulin tolerance tests, mice were fasted for 6 h and then i.p. injected with insulin (0.8-1.2 units/kg body weight). Blood glucose concentrations were measured before and 30, 60, 90, and 120 min after insulin injection.

### Hyperinsulinemic euglycemic Clamps

Mice with an inserted catheter into the right internal jugular vein were prepared as previously described [36]. Mice were equilibrated from t = -90 to 0 min after 4–6 h fasting. [3-^3^H] glucose (3 mCi; Moravek, California, USA) was administered at t = -90 min, followed by a constant infusion of 0.05 μCi/min. After 90 min as a basal period (t = -90 to 0 min), blood samples were collected from the tail vein for the determination of plasma glucose concentration and basal glucose-specific activity. Then the continuous human insulin (Humulin; Novo Nordisk) infusion was started (t = 0 min) at a rate of 4 mU/kg/min to keep hyperinsulinemic condition with submaximal suppression of HGP to assess insulin sensitivity. At 0 min, the continuous infusion rate of [3-^3^H]-D-glucose tracer was increased to 0.15 μCi/min for minimization of the changes in glucose-specific activity. At t = 75 min, 10 μCi 2-[^14^C] D-glucose (Moravek, California, USA) was administered to each mouse to measure glucose uptake. During the 120 min clamp, variant glucose was simultaneously infused to keep the blood glucose concentration stable (∼130 mg/dL). At the end of the clamp, liver, muscle, and adipose tissues were collected for the determination of radioactivity. Serum and tissue radioactivity were measured and calculated as previously described [36].

### Insulin sensitivity evaluation of liver

Tissue insulin action was evaluated by measuring insulin-stimulated AKT phosphorylation in the liver. Briefly, mice were i.p. injected with 2 units/kg insulin for 10 min injection or saline after 5 h of fasting, then sacrificed for liver collection to measure the phosphorylation of AKT and total AKT.

### Histological analysis

For H&E staining, tissues were embedded in paraffin and sectioned at 5 μm thickness. For Oil Red O staining, livers were embedded in Tissue Freezing Medium (Leica, Germany) and sectioned to 15 μm using a Leica Cryostat. For lipid size quantification, lipid drop diameters were measured in the H&E-stained sections of three individual samples in each group using ImageJ.

### Hepatocytes Oil-red O staining

Wash hepatocytes with PBS twice and then fix cells with 3.7% formaldehyde solution for 20 min. Then cells were washed with PBS twice. Then incubate fixed hepatocytes with Oil-red O working solution (0.5% Oil-red O: ddH_2_O = 3:2) for 30 min. Wash cells with ddH2O several times and the stained fat droplets in the cells were visualized by light microscopy and photographed. Add 200-500 μL isopropanol to each well, gently mix for 15 min, and read the absorbance at 490 nm with a 96-well plate reader to quantify the accumulation of lipids.

### Quantification of liver TG in patient

The patients underwent CT examination in a supine position with a multidetector spiral CT scanner (VCT, GE Healthcare, Milwaukee, WI, USA). The scan ranged from the diaphragm to the pubic symphysis. CT images were obtained with a tube voltage of 120 kV, a tube current of 240 mA, a slice thickness of 5 mm, a slice interval of 5 mm, a rotation time of 0.6 s, a helical pitch of 1.375, a field of view of 35–40 cm, a matrix of 512 × 512, and a standard reconstruction algorithm.

### Hepatocytes lipid droplet staining

Hepatocytes were stained with Lipid Droplets Fluorescence Assay Kit (BioVision, USA) according to the manufacturer’s instructions. Examine lipid droplet staining with a fluorescence microscope equipped with a filter designed to detect FITC (excitation/emission=485/535nm). Cells were prepared according to the manufacturer’s instructions to measure the lipid droplet-stained positive cells.

### Immunofluorescence microscopy

Cells (1 × 10^5^) were seeded on glass coverslips for 18 h, fixed in 4% paraformaldehyde, quenched with 0.1 M glycine, permeabilized in 0.2% BSA-0.05% saponin in PBS, and incubated with primary antibodies targeting Perilipin4, Bodipy (Sigma, USA) followed by treatment with fluorescent-labeled secondary antibody and DAPI (Santa Cruz Biotechnology, USA). Images were acquired on a Two-Photon Laser Confocal Microscope (Leica TCS SP8-MP).

### Subcellular fractionation

Subcellular primary hepatocyte fractionation was performed using the Triton X-114 partition method and ultracentrifugation [37]. In brief, hepatocytes with the indicated treatment were lysed in 500 μl lysis buffer. The total protein concentration was diluted to 1 mg/ml and partitioned with the same volume of 4% Triton X-114 for 5 min at 37 °C to solubilize and fractionate the lipid-rich cell membrane. The aqueous upper phase contains enriched intracellular protein, and the organic lower phase contains highly enriched membrane-associated proteins. All the samples were subjected to immunoprecipitation and western blot detection of the proteins that were present at different portions.

The culture flask was filled with DMEM/F12 medium containing 20% fresh calf serum and placed upside down in a 37°C, 5% CO2 satiated incubator. The cells firmly adhered to the inner surface of the culture flask. After the cells firmly adhered to the inner surface of the culture flask, the flask was placed normally for incubation

### RNA isolation, Reverse transcription PCR, and Real-time PCR

RNA from tissues was isolated using TRIzol reagent (Takara Bio, Japan) and reverse transcribed using PrimeScript RT Master Mix for RT-PCR (Takara Bio, Japan). Real-time PCR was performed using SYBR Select Master Mix (Applied Biosystems, USA) on an ABI 7300 sequence detector (Applied Biosystems, USA). For RT-PCR, cDNA was synthesized using Taq-Man RNA reverse transcription kit and RNA primers (5x). qPCR was performed using TaqMan universal master mix II and RNA primers (20x) in 10 μL reactions on a StepOnePlus Real-Time PCR Systems (ThermoFisher Scientific). The results were normalized to the 18S level in each sample, and endogenous U6 small nuclear RNA was used for the normalization of cellular or plasma RNA. All reactions were carried out in triplicate, and analysis was carried out using the 2^−ΔΔCt^ method.

### Glucose uptake assay

After 8 h serum starvation, cells were stimulated with 100 nM insulin for 30 min in KRH buffer (137 nM NaCl, 4.8 mM KCl, 1.2 mM KH_2_PO_4_, 1.2 mM MgSO_4_, 2.5 mM CaCl_2_, 0.2% BSA, 16 mM HEPES) at 37℃. Then 3H-2-deoxy-D-glucose (3H-2-DOG, 0.1 mM, 0.4 μCi/mL) was supplemented to cells. After 10 min incubation at 37℃, cells were washed with ice-cold PBS twice. NaOH (1 N) was then added and incubated for 20 min to efficiently dissolve cells. An aliquot was used for protein concentration measurement. After neutralizing NaOH by adding HCl (1 N), the extracts were transferred to scintillation vials, scintillation fluid was added and the radioactivity was counted. Results were normalized with the protein concentration of cell lysates.

### Glucose output assay

After 6 h of serum starvation, the primary hepatocytes were washed twice and then exposed to glucose-free buffer (10 mM HEPES, 4 mM KCl, 125 mM NaCl, 0.85 mM KH_2_PO_4_, 1.25 mM Na_2_HPO_4_, 1 mM CaCl_2_, and 15 mM NaHCO_3_) containing glucagon (200 ng/mL), insulin (10 nM), or a combination of glucagon and insulin for 4 h, at 37℃. Glucose production was determined by the measurement of glucose in the media. The primary hepatocytes attached to the culture plate were dissolved by adding NaOH (1 N) and protein content was determined. The glucose results were normalized with the protein concentration of cell lysates.

### Protein extraction and Western Blot Analysis

Cells or tissues were homogenized in RIPA buffer supplemented with protease inhibitors and phosphatase inhibitor cocktails (Roche Diagnostics, Indianapolis, IN, USA). Western blot analysis was performed according to the routine procedure. Proteins were quantified with the BCA method. Approximately 30-50 μg of proteins were separated by 10% SDS-PAGE and transferred onto a PVDF membrane (Roche). The membrane was blocked with 5% non-fat milk in PBS for 1 h at room temperature and then incubated with the specific primary antibodies at 4°C overnight. The bonded antibodies were detected by the secondary antibodies, diluted at 1:10000 in PBS, for 1 h at room temperature. The signals were detected with an enhanced chemiluminescence system. Western blot data in figures and supplemental figures are all representatives of more than three independent experiments.

## QUANTIFICATION AND STATISTICAL ANALYSIS

Blinding was performed whenever deemed to be appropriate and applicable. During data plotting and analyses, sample description and identification were unavailable to the core personnel. No samples or data were excluded from the study for statistical purposes. Each *in vitro* experiment was independently performed with duplicate or triplicate to ensure reproducibility. Group sizes of 6 mice or above were sufficient to reach a statistical power of at least 80%. Mice were assigned at random to treatment groups for all mouse studies. Tests used for statistical analyses are described in the figure legends. The sample numbers are mentioned in the figure legends. To assess whether the means of the two groups are statistically different from each other, an unpaired two-tailed Student’s t-test was used for statistical analyses, all data passed the normality test using Prism8 software (GraphPad software v8.0; Prism, La Jolla, CA). p values of 0.05 or less were considered to be statistically significant. Degrees of significance were indicated in the figure legends. For glucose and insulin tolerance test results, statistical comparisons between every two groups at each time point were performed with an unpaired two-tailed Student’s t-test.

## Author Contributions

Conceptualization, C.J.L., X.H and Y.B.; Methodology, X.H., and L.F.; Formal Analysis: Y.Z., and H.Y.N.; Investigation, Y.Z., H.Y.N., M.F.Z., P.S., S.J., J.Z.Z., X.C.W., and Y.P.T.; Resources, X.W.Y, X.T.S, X.D.S., L.F., and J.H.L.; Writing – Original Draft, Y.Z., and H.Y.N.; Writing – Review & Editing, C.J.L., Y.Z., Y.B., X.H., and L.F.; Funding Acquisition, Y.Z., C.J.L., L.F.; Supervision, C.J.L.

## Acknowledgments

This work was supported by Grants from the National Natural Science Foundation of China (32271182) to Yue Zhao, the Outstanding Youth Foundation of Jiangsu Province of China (BK20200061) to Yue Zhao, the National Natural Science Foundation of China (31770838) to Lei Fang, and the National Natural Science Foundation of China (91857109) to Chaojun Li.

---

**Figure S1.**
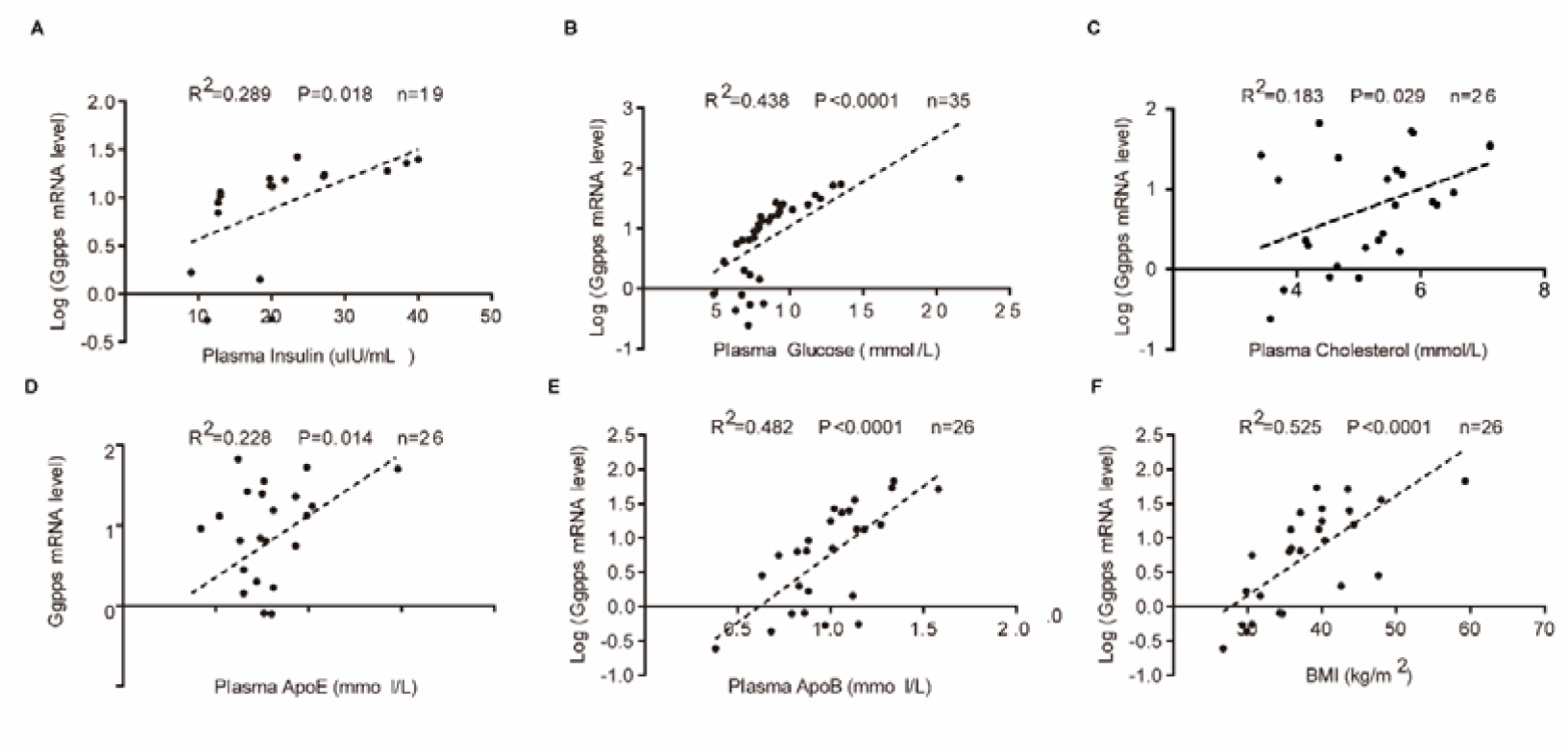
Related to Figure 1. Characteristics of MHO and MAO individuals. Linear regression analysis between plasma insulin (A), plasma glucose (B), plasma cholesterol (C), plasma ApoE (D), plasma ApoB (E), BMI (F), and *Ggpps* mRNA levels in patients (n = 19-35). All experiments were repeated at least twice with similar results. *p < 0.05, **p < 0.01, ***p < 0.001. Data are represented as mean ± SEM. Two-sided Student’s t-test.

**Figure S2.**
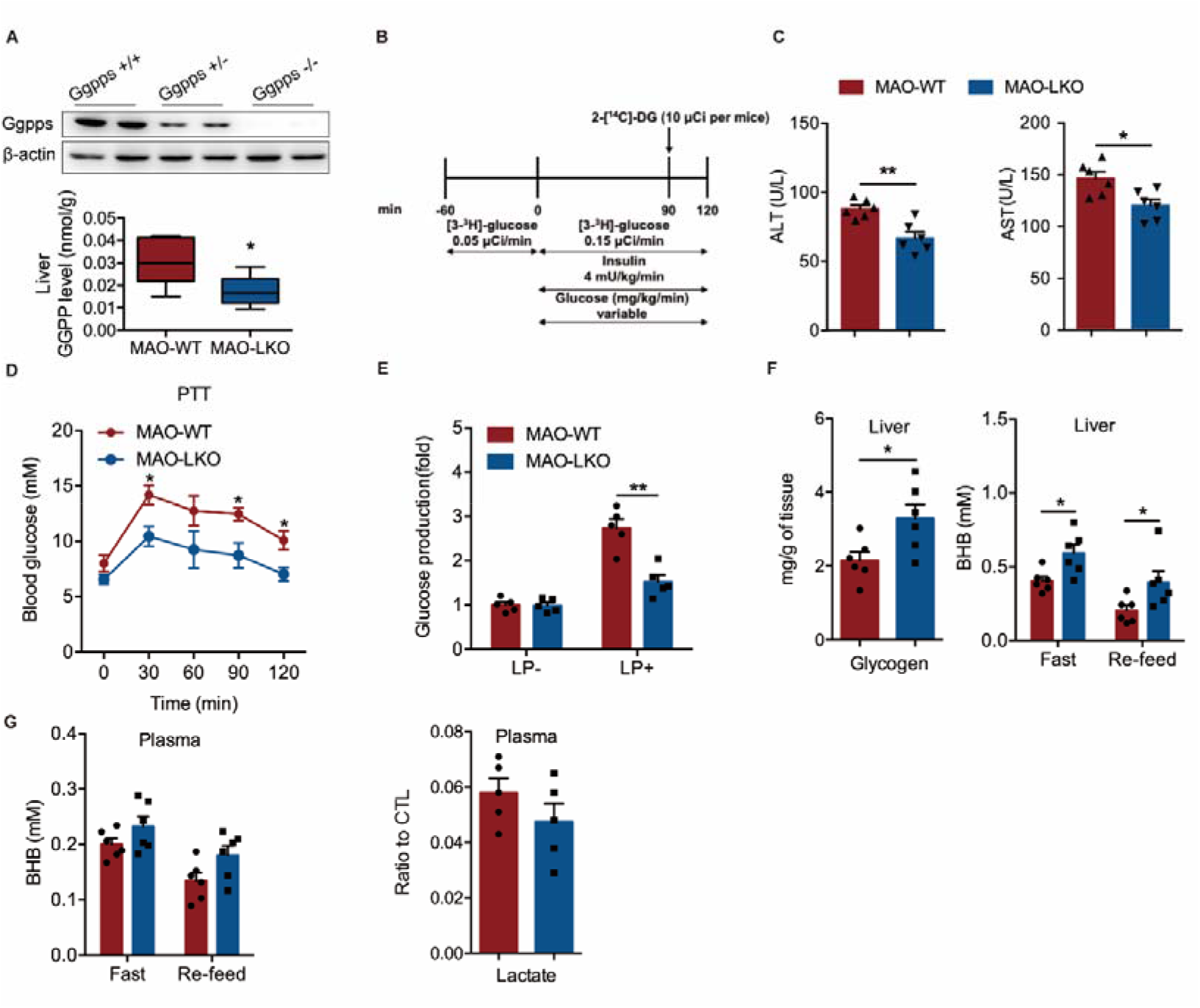
Related to Figure 2. The effect of GGPP on liver metabolism of MHO and MAO. (A) Western blots analysis of Ggpps expression and the liver GGPP level (n = 6) in wild-type (WT) and L-*Ggpps* KO mouse livers. (B) Schematic representation of the experimental procedure of hyperinsulinemic-euglycemic clamp assay in MAO-WT and MAO-LKO mice. (C) ALT and AST in the liver of WT and LKO mice under MHO or MAO conditions. (n = 6). (D) Pyruvate tolerance tests in MAO-WT and MAO-LKO mice. (n = 8). (E) Glucose production in primary hepatocytes of MAO-WT and MAO-LKO mice after lactate and pyruvate (LP) treatment. (n = 5). (F) Liver glycogen, β-hydroxybutyrate (BHB) in overnight-fasted or re-feed MAO-WT and MAO-LKO mice. (n = 6). (G) Plasma BHB and lactate in MAO-WT and MAO-LKO mice. (n = 5 or 6). All experiments were repeated at least twice with similar results. *p < 0.05, **p < 0.01, ***p < 0.001. Data are represented as mean ± SEM. Two-sided Student’s t-test.

**Figure S3.**
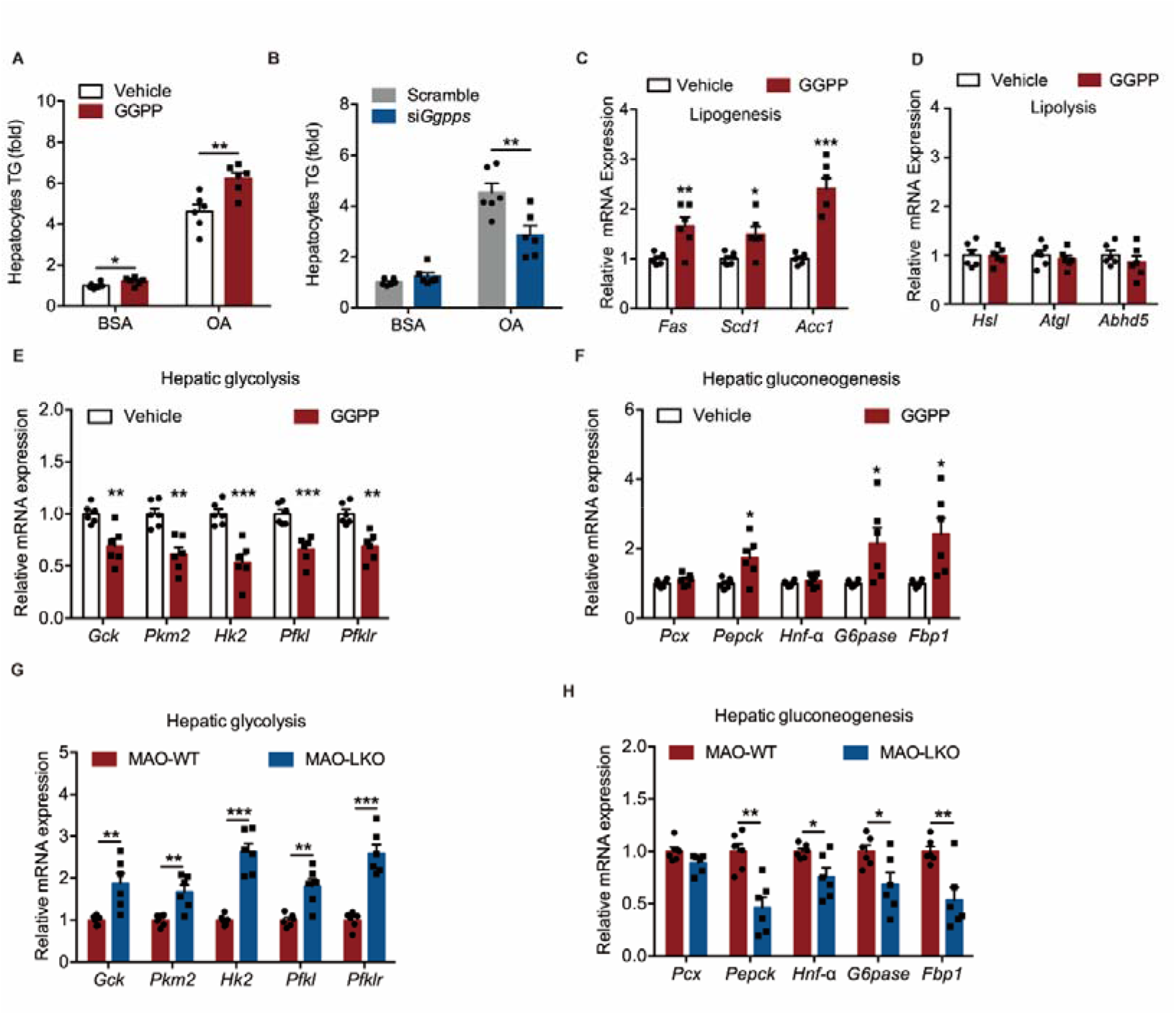
Related to Figure 3. Characteristics of the liver in WT and LKO mice under MAO condition. (A) TG quantification of primary hepatocytes treated with vehicle or GGPP under BSA or OA treatment. (B) TG quantification of primary hepatocytes treated with scramble or si*Ggpps* under BSA or OA treatment. (C-D) Expression of genes related to lipogenesis and lipolysis in the hepatocytes treated with GGPP. (E-F) Expression of genes related to glycolysis (E) and gluconeogenesis (F) in the primary hepatocytes treated with vehicle or GGPP. (n = 6). (G-H) Expression of genes related to glycolysis (G) and gluconeogenesis (H) in the liver of MAO-WT and MAO-LKO mice. (n = 6). All experiments were repeated at least twice with similar results. *p < 0.05, **p < 0.01, ***p < 0.001. Data are represented as mean ± SEM. Two-sided Student’s t-test.

**Figure S4.**
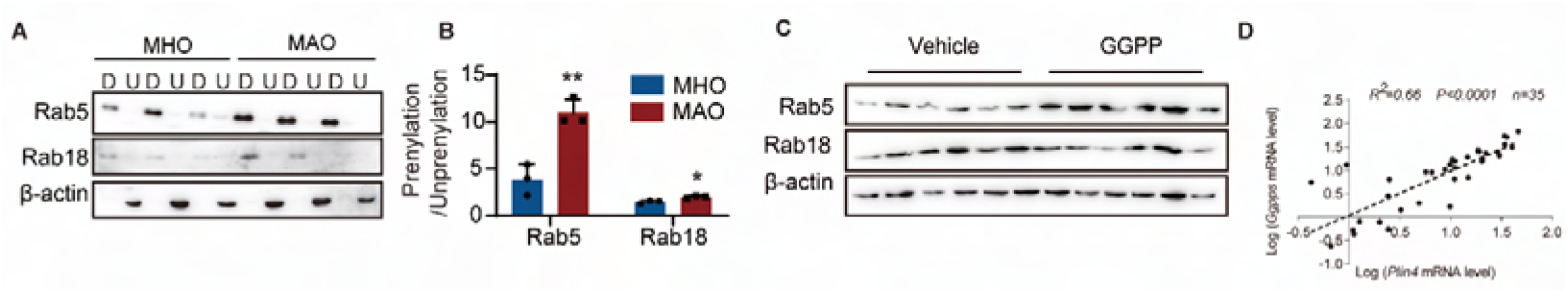
Related to Figure 4. The effect of GGPP-dependent Perilipin4 prenylation on insulin sensitivity *in vivo*. (A-B) Membrane-associated Rab5 and Rab18 (lipid-soluble protein with prenylation in the detergent phase, down) and cytoplasm-associated Rab5 and Rab18 (water-soluble protein with no prenylation in the aqueous phase, up) were obtained by Triton X-114 extraction and analyzed by immunoblotting. (n =3). (C) Rab5 and Rab18 protein levels in primary hepatocytes following GGPP treatment. (D) Linear regression analysis between *Perilipin4* mRNA levels and *Ggpps* mRNA levels in patients. (n = 35). All experiments were repeated at least twice with similar results. *p < 0.05, **p < 0.01, ***p < 0.001, *n.s.*, no significant difference. Data are represented as mean ± SEM. Two-sided Student’s t-test.

**Figure S5.**
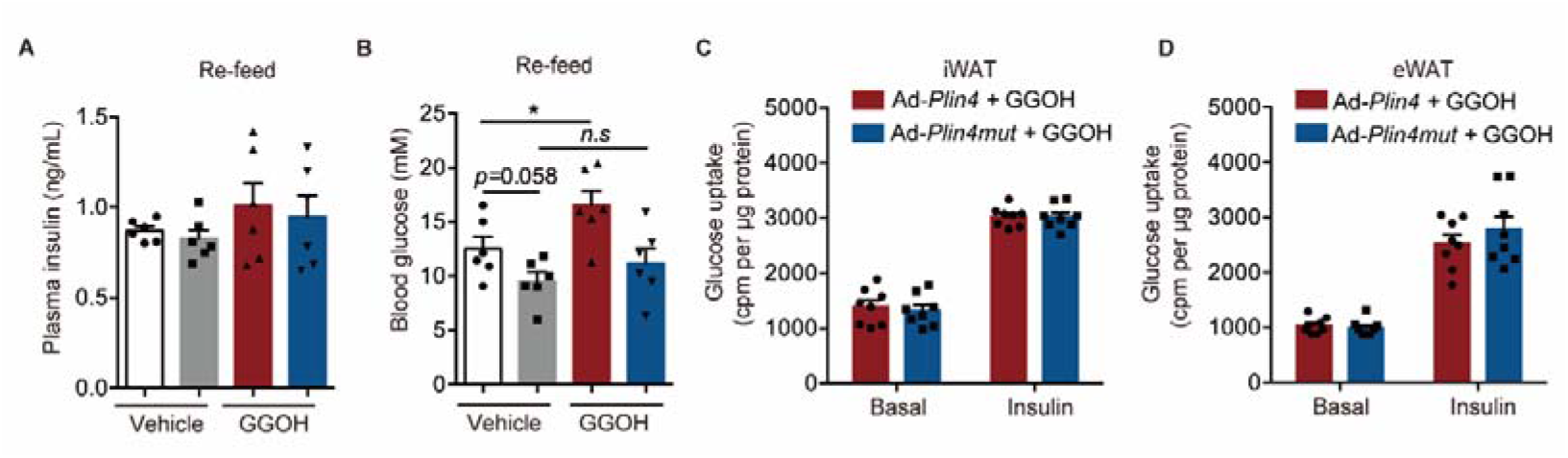
Related to Figure 5. Plin4C1500S inhibits GGPP-induced hepatic lipid droplet formation and ameliorates GGPP-induced insulin resistance. (A-B) Insulin level and Blood glucose in overnight-fasted and re-feed mice (n = 6). (C-D) Glucose uptake in primary adipocyte of iWAT (H), eWAT (J) from E. (n = 8). All experiments were repeated at least twice with similar results. *p < 0.05, **p < 0.01, ***p < 0.001, *n.s.*, no significant difference. Data are represented as mean ± SEM. Two-sided Student’s t-test.

**Figure S6.**
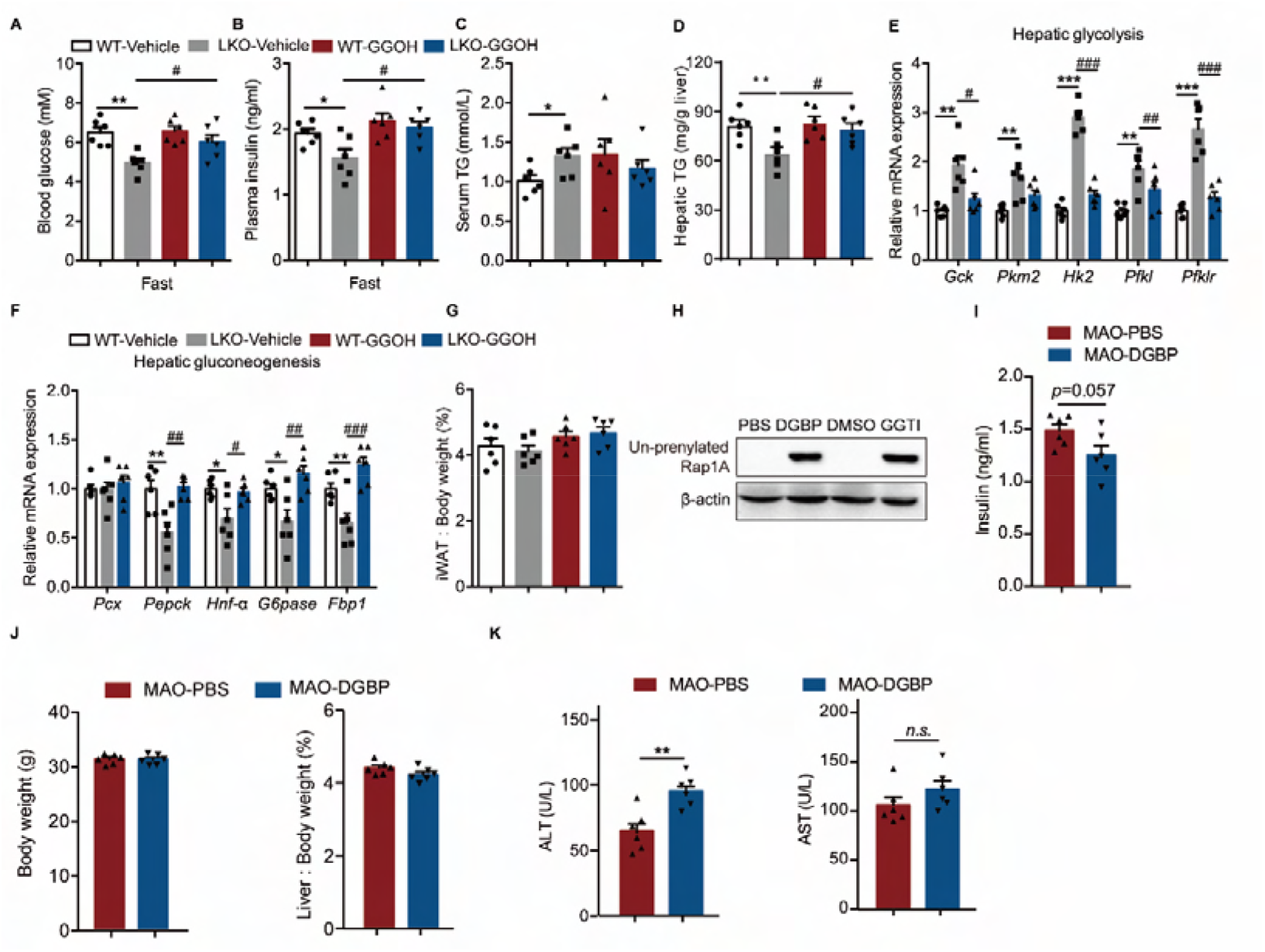
Related to Figure 6. The effect of Ggpps inhibitor on metabolic phenotypes. (A-D) Blood glucose (A), blood insulin (B), serum TG (C), and hepatic TG(D) in WT or LKO mice with vehicle or GGOH treatment. (n = 6). (E-F) Expression of genes related to glycolysis (E) and gluconeogenesis (F) in the liver of WT or LKO mice with vehicle or GGOH treatment. (n = 6). (G) iWAT weight relative to the whole-body weight of mice from WT or LKO mice with vehicle or GGOH treatment. (n = 6). (H) Western blots of Un-prenylated Rap1A in primary hepatocytes treated with Ggpps inhibitor-DGBP and GGTases inhibitor (GGTI). (I) Plasma insulin in MHO and MAO mice with PBS or DGBP treatment. (J) Body weight and liver weight relative to the whole-body weight of mice. (K) ALT and AST in MAO mice with PBS or DGBP treatment. (n = 6). All experiments were repeated at least twice with similar results. *p < 0.05, **p < 0.01, ^#^p < 0.05, ^##^p < 0.01, ^###^p < 0.001. *n.s.*, no significant difference. Data are represented as mean ± SEM. Two-sided Student’s t-test.

